# Rapid replacement of established by exotic genetic lineages of the fungal maize pathogen *Exserohilum turcicum* in the Swiss Rhine valley

**DOI:** 10.1101/2025.06.24.661258

**Authors:** Mireia Vidal-Villarejo, Michael Hammerschmidt, Barbara Oppliger, Hans Oppliger, Karl Schmid

## Abstract

The fungal maize pathogen *Exserohilum turcicum*, the causative agent of Northern Corn Leaf Blight (NCLB) was introduced to Europe in the late 19th century, *E. turcicum* where it rapidly expanded. It persisted in four major clonal lineages and multiple physiological races, defined by their interactions with maize disease resistance genes. In a metagenomic survey of natural infections on the susceptible traditional landrace ‘Rheintaler Ribelmais’ in the Swiss Rhine Valley during 2016 and 2017, we found that one of these clonal lineages (‘Small clonal’ cluster) was dominant in the region. We repeated the surveys in subsequent years and applied a novel pooling strategy to facilitate large-scale sampling by combining ten infected leaves per sampling location. This approach increased sample throughput while reducing sequencing and laboratory costs. The new survey revealed in 2021 and 2022 a significant temporal shift in population structure of exotic genetic lineages from the tropical ‘Kenyan’ cluster, which have become predominant. It indicates recent introduction and establishment of genetically diverse, tropical *E. turcicum* lineages into a temperate agricultural system, possibly facilitated by climate change and global seed exchange. Phyllobiome analyses of infected leaves showed that microbial community composition varied across years but remained largely consistent between maize variety types (landrace vs. hybrid). Overall, metagenomic pool sequencing of infected leaves proves to be a cost-effective, spatially resolved method for pathogen monitoring and provides useful evidence for the evolving epidemiology of *E. turcicum*, with implications for developing durable resistance in maize breeding programs.

## Introduction

Climate change and human activities, such as the globalization of agricultural systems, have contributed to the spread of plant pathogens (***Bebber et al., 2013; Ristaino et al., 2021; Chaloner et al., 2021; Miedaner and Garbelotto, 2024***). A fundamental characteristic of plant pathogen epidemics is the rapid evolution and demographic changes of plant pathogens, because both plants and their pathogens are constantly evolving and co-adapting (***Clay and Kover, 1996***). Consequently, close monitoring of pathogen epidemiology is essential to remain competitive in this evolutionary arms race (***Jeger et al., 2023***). Monitoring and prediction systems are being developed, incorporating climate change models to predict pathogen pressure and provide decision support for farmers (***Ristaino et al., 2021; Singh et al., 2023***). Traditional approaches to identifying novel pathogen strains, such as phenotyping with differentiation panels of plant genotypes carrying different resistance genes, are slow and and make rapid response difficult.

The advancement of DNA sequencing techniques offers new possibilities for characterizing the spatiotemporal dynamics of pathogen epidemiology (***Weisberg et al., 2021***). Although predicting the pathogenicity of various pathogen genotypes and understanding plant-pathogen interactions for effective mitigation remains challenging (***Jones et al., 2018; Barwell et al., 2021; Seong and Krasileva, 2023***), monitoring based on DNA sequencing could guide crop rotation strategies to reduce pathogen population sizes, predict the geographic spread of pathogens, and inform variety selection and resistance breeding programs (***Wang et al., 2021; McDonald et al., 2022; Wuest et al., 2021***). DNA sequencing may uncover novel virulence genes and interactions between plants, pathogens, and the microbiome, which could subsequently benefit agricultural practices (***Jones et al., 2018; Feurtey et al., 2023; Vidal-Villarejo et al., 2024***). Large reference collections of pathogen strains support the modeling of pathogen dynamics and the identification of evolutionary adaptation patterns to develop robust plant protection strategies (***Feurtey et al., 2023; Vidal-Villarejo et al., 2023***).

Northern Corn Leaf Blight (NCLB), which is caused by the fungal pathogen *Exserohilum turcicum* (syn. *Setosphaeria turcica*) is major disease of maize, because it it occurs globally and can cause substantial yield losses (***Galiano-Carneiro and Miedaner, 2017***). Consequently, resistance to this pathogen has been a long-standing focus of maize breeding efforts, and the ongoing climate change maintains this requirement (***Miedaner and Juroszek, 2021***). The diversity of *E. turcicum* has been studied extensively over the past decades (***Borchardt et al., 1998c***,***Borchardt et al., 1998a***; ***Galiano-Carneiro and Miedaner, 2017; Vidal-Villarejo et al., 2023***). These studies revealed that *E. turcicum* reproduces sexually in tropical regions, whereas clonal reproduction predominates in temperate regions, leading to reduced genetic diversity in those areas. Differential screening has identified pathogen races based on their ability to infect maize varieties with different resistance genes (***Hanekamp, 2016***). However, predicting the virulence phenotype of an *E. turcicum* strain based solely on its DNA sequence remains challenging (***Vidal-Villarejo et al., 2023***).

In earlier work, we characterized the genetic diversity of *E. turcicum* through whole-genome resequencing of isolates collected across Europe and northeastern Africa and identified a small number of dominant clonal lineages in Europe (***Vidal-Villarejo et al., 2023***). Despite the low genetic diversity at the lineage level, we found substantial genetic diversity of effectors and potentially other virulence-related genes within these lineages, suggesting that these genetic variants likely influence the ability of specific isolates to infect different host plants. Our previous monitoring study conducted in the Swiss Rhine Valley (2016–2018) using metagenomic sequencing of infected leaves from the susceptible landrace ‘Rheintaler Ribelmais’ documented the presence of all major *E. turcicum* clonal lineages in this region and suggested that pathogen diversity originated from local soil inoculum rather than seed stock (***Vidal-Villarejo et al., 2024***). While microbiome composition showed significant associations with pathogen DNA abundance, the specific clonal lineage of *E. turcicum* had minimal impact on overall microbiome diversity.

Here, we extend our pathogen monitoring in the Swiss Rhine Valley by including four additional years of data and adopting a pooled sampling strategy, in which we combined multiple leaf samples from individual fields to reduce sampling and preparation costs. Despite lower costs, metagenomic sequencing retains high resolution, enabling the identification of distinct *E. turcicum* clonal lineages within pooled samples. Our monitoring includes both the highly susceptible traditional cultivar ‘Rheintaler Ribelmais’, which is used as a sentinel plant for early detection of invasive strains (***Lovell-Read et al., 2023***), and modern maize hybrids bred for NCLB resistance. We integrate this new dataset with previous metagenomic surveys and whole-genome sequences from our reference collection to demonstrate the effectiveness of pooled metagenomic sequencing for pathogen surveillance. The results reveal a recent increase in genetic diversity, driven by the introduction of novel exotic, tropical lineages, suggesting the influence of climate change and emphasizing the need for adaptive plant protection strategies in European agriculture.

## Results

### Sampling and metagenomic pool sequencing

Leaves from maize plants naturally infected with NCLB were collected from the suceptible traditional cultivar ‘Rheintaler Ribelmais’ and commercial hybrid varieties, whose identity was not recorded. The sampling was conducted on agricultural production fields at a total of 13 distinct locations in the Swiss Rhine Valley between the years 2019 and 2022 (Supplementary Dataset S1). We sequenced 60 pooled samples using a shotgun metagenomics strategy which are sum-

Each sequencing sample consists of a pool of approximately 10 leaves from the same field. For comparison with the pooled samples, we also sequenced three single leaves of three collection sites from 2021. On average, 10.7 Gbp of sequencing data per sample was obtained and a mean of 72.1% of the reads mapped to the maize reference genome. The taxonomic classification with Kaiju revealed that nearly all (19 out of 20) samples from 2021 have a high percentage of reads classified as *Exerohilum*, in contrast, samples from 2019, 2020, and 2022 *Exerohilum* reads are scarce, and instead have a high percentage of reads classified as the common rust on maize (genus *Puccinia*; Fig. 1). The dominant bacterial taxa are consistent across all samples, except in cases with a high presence of *Exserohilum* reads (almost all samples from 2021 and few from 2022), which have an apparent increase in the abundance of the genus *Klebsiella* compared to samples with no or little *Exserohilum* DNA.

**Figure 1.**
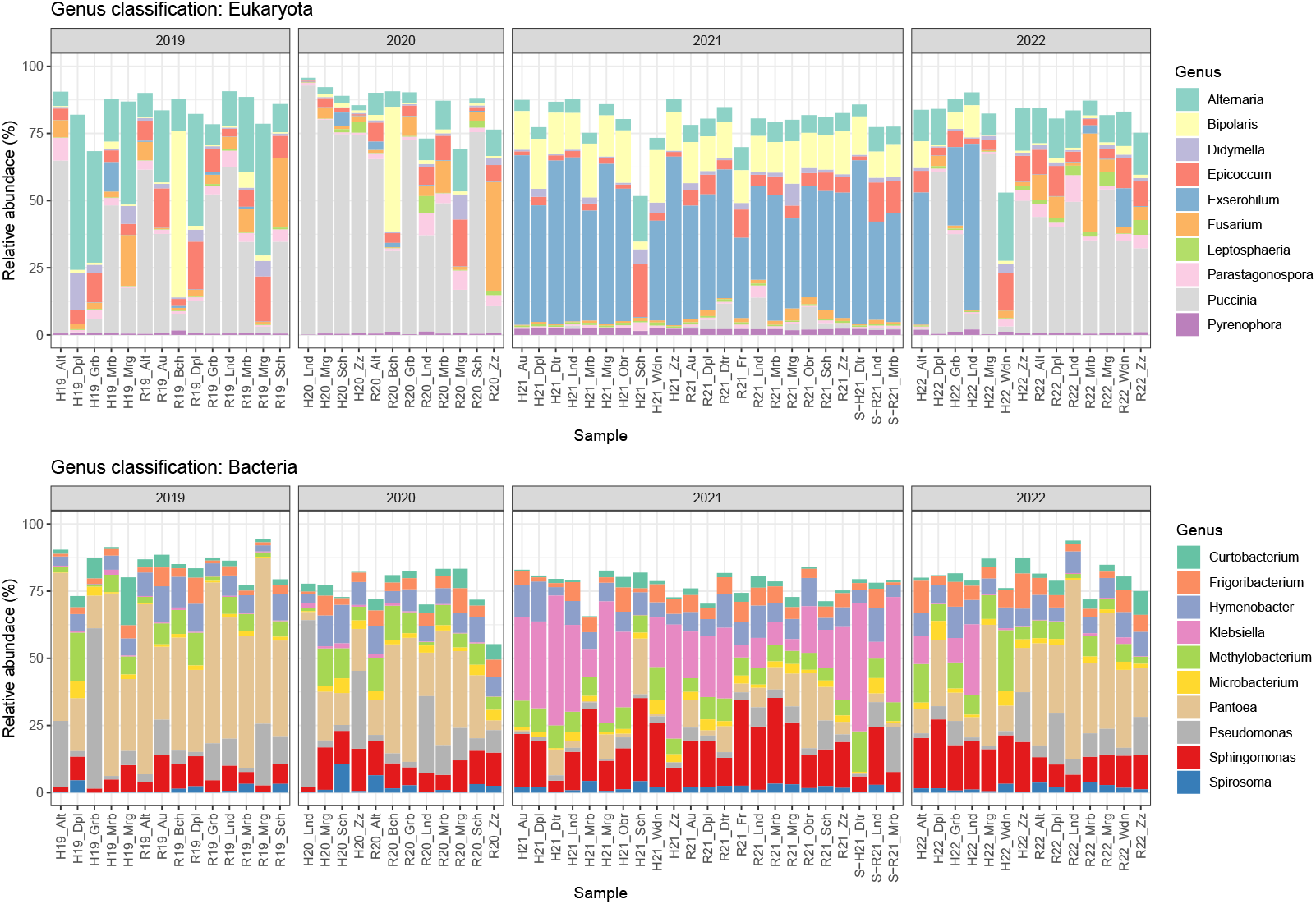
Relative abundance of the top 10 most abundant taxons (at genus level) in Eukarya (top) and Bacteria (bottom). Percentage relative to the total number of reads classified as Eukaryotic or Bacterial.

### Metagenomic classification of *E. turcicum* and variant filtering

Reads classified as genus *Exserohilum* were mapped to the reference genome of *E. turcicum* (*Setosphaeria turcica* Et28A v1.0; ***Ohm et al., 2012; Condon et al., 2013***) followed by single nucleotide polymorphism (SNP) calling. A median of 0.08% of the total non-host metagenomic reads mapped to the *E. turcicum* reference genome, with a mean coverage of 0.15X. Consistent with Kaiju classification results (Fig. 1), a markedly higher proportion of reads from the 2021 samples mapped to *E. turcicum* (median: 10.85%), with a mean coverage of 23X (Supplementary Fig. S1). SNP calls from these samples were merged with a VCF file containing variant data from 121 isolates from Europe and Kenya with whole-genome sequencing (hereafter WGRS isolates) (***Vidal-Villarejo et al., 2023***), as well as metagenomic samples from the Swiss Rhine Valley collected in 2016 and 2017 and a previous pilot study (***Vidal-Villarejo et al., 2024***). After merging, we retained only biallelic SNPs and excluded positions with more than 40% missing data, resulting in a combined dataset of 10,260 SNPs. Further filtering excluded positions with fewer than 2 reads or with excessively high coverage (above the 90th percentile), treating them as missing. Subsequently, we removed all positions with more than 20% missing data and samples with more than 50% missing loci. The final filtered SNP set contained 7,750 SNPs across 181 samples, including 22 from 2021 and four from 2022. Due to the low number of *Exserohilum*-classified reads and subsequent stringent filtering, no samples from 2019 or 2020 were retained in the final SNP set. To recover representative data from these years, we applied a modified filtering workflow that allowed us to retain two samples from 2019 (H19_Mrb and R19_Grb) and two from 2020 (H20_Sch and R20_Alt) based on the highest number of non-missing positions. These recovered datasets were derived from the soft-filtered SNP set of 10,260 SNPs by retaining only those positions without missing data in the low-coverage sample, and applying a threshold of 20% missing data per position and 50% per sample afterwards. The recovered datasets retained 732 and 64 positions for the 2019 samples, and 216 and 181 positions for the 2020 samples, respectively. Using the same method, we also recovered one additional 2022 sample (R22_Grb), which retained 1,551 SNPs.

### Inference of *E. turcicum* genetic structure

We previously described five genetic clusters of *E. turcicum*: four from Europe and one from Kenya (***Vidal-Villarejo et al., 2023***). Three of the five clusters correspond to European clonal lineages, which we named ‘Big Clonal’, ‘Small Clonal’, and ‘French Clonal’. These lineages are characterized by low genetic diversity, the presence of only one mating type, and absence of recombination. The remaining two clusters, named ‘Diverse’ and ‘Kenyan’ lineages, represent genetically diverse populations with high genetic variability, the presence of both mating types, and evidence of past sexual recombination. To infer population structure of all available samples we first constructed a neighbor-joining (NJ) tree based on allele frequencies derived from read counts (Fig. 2A). In the resulting tree most samples from 2021 and 2022 form a distinct cluster, positioned between the Kenyan-French Clonal and Diverse lineages.

**Figure 2.**
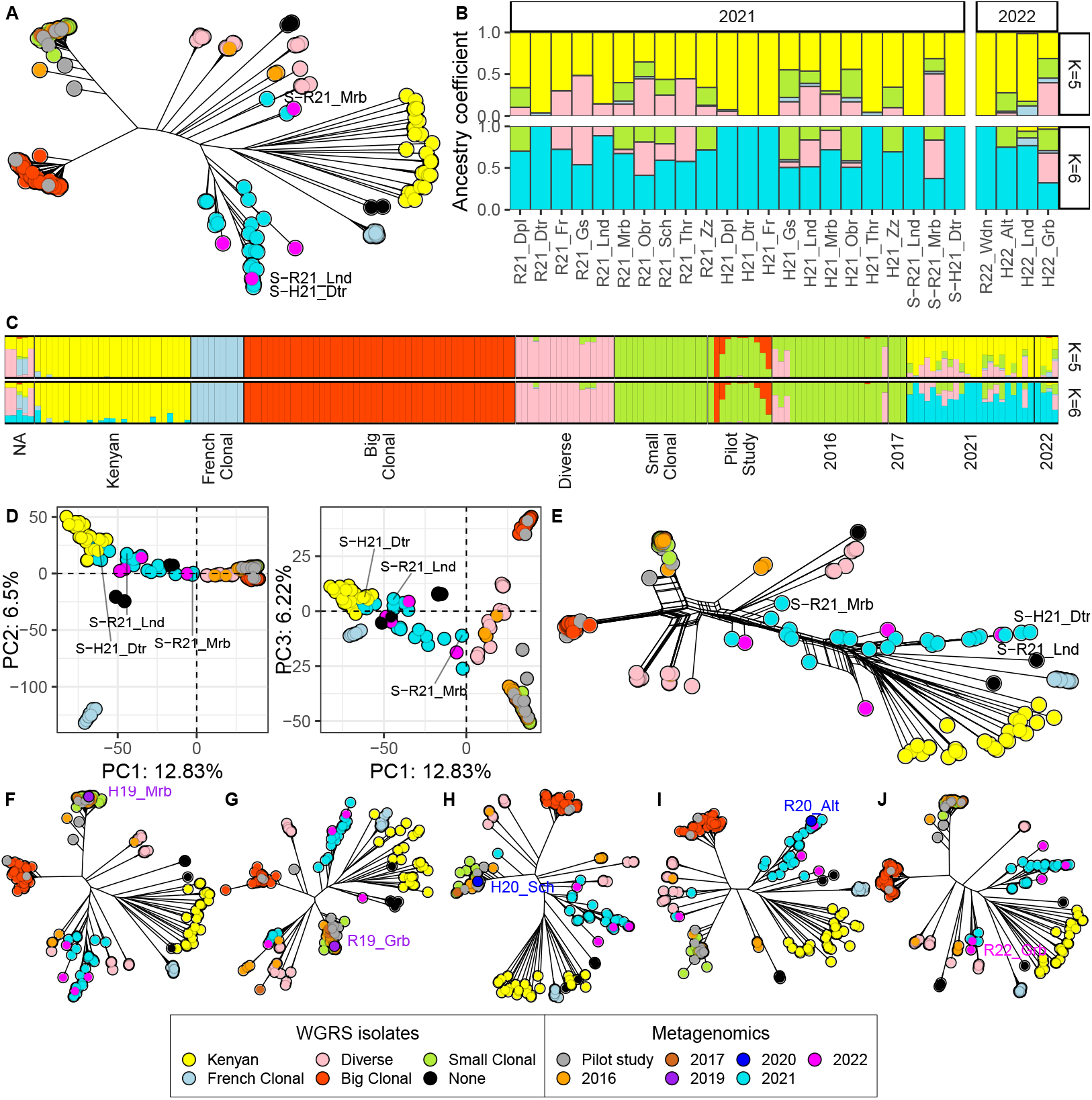
**A)** NJ tree with European and Kenyan WGRS isolates and metagenomic samples from 2016, 2017, 2021, 2022 and the pilot study. The three non-pooled samples from 2021 are indicated with their name. **B)** Ancestry coefficients for K=5 and K=6 of the metagenomic samples from 2021 and 2022 **C)** Ancestry coefficients for K=5 and K=6 **D)** PCA and **E)** Neighbour-Net of the metagenomic samples from 2016, 2017, 2021, 2022 and European and Kenyan WGRS isolates. Axis in the PCA indicate the percentage of variance explained by each component. **F-J)** NJ trees of five low-covered recovered samples from (**F-G**) 2019, (**H-I**) 2020 and (**J**) 2022. Name of the recovered samples are shown in the trees. WGRS isolate samples are colored according to the five defined European and Kenyan lineages and metagenomic samples are colored according to the year of collection.

We then inferred ancestry coefficients with ADMIXTURE. Consistent with the NJ tree, the new samples from 2021 and 2022 predominantly exhibit ancestry from the Kenyan lineage, with contributions from the Diverse and Small Clonal clusters (Fig. 2B and C). With *K* = 6 (and higher) the 2021 and 2022 samples have a high degree of ancestry of a new, distinct genetic group, mirroring the structure observed in the NJ tree (Fig. 2B-C and Supplementary Figure S2). A total of eight samples from 2021 and 2022 are exclusively assigned to this new genetic group (Fig. 2B). Results from a principal Component Analysis (PCA) and a Neighbor-Net (NNet) analysis partially differ from the previous analyses. In the PCA, the 2021 and 2022 samples do not form a distinct cluster, but are spread between the Kenyan and Diverse lineages or cluster with the Kenyan lineage (Fig. 2D). In the NNet analysis, some samples also appear admixed between the Diverse and Kenyan lineages, as indicated by the reticulate network, whereas other samples form a distinct cluster (Fig. 2E), a pattern consistent with the NJ tree (Fig. 2A).

Despite the low SNP counts, NJ trees, PCA, and NeighborNet analyses of low-coverage samples from 2019–2022 consistently placed them within established *E. turcicum* lineages: Three samples (two 2019 samples and one 2020) within the Small Clonal cluster; one 2020 sample with the cluster with most of the 2021 samples; and the 2022 sample within a Diverse lineage (Figs. 2F–J, Supplementary Fig. S3).

Despite the limited number of SNPs in these recovered samples, the structure of the five *E. turcicum* lineages is generally maintained.

### Pooled leaves vs. single-leaf samples

To clarify whether the apparent new genetic cluster in 2021–2022 represents a truly novel *E. turcicum* lineage or an artifact of pooled sequencing, we reanalyzed population structure using NJ trees, PCA, NNet, and ADMIXTURE with pooled samples excluded. We retained only the three non-pooled 2021 samples along with single-leaf metagenomic samples from 2016 and 2017 and the pilot study to test whether the previously observed cluster persisted without pooled data. The NJ tree, NNet, PCA, and ADMIXTURE analyses show that two of the samples (S-R21_Lnd and S-H21_Dtr) cluster with the Kenyan lineage and exhibit corresponding ancestry coefficients, while the third sample (S-R21_Mrb) clusters with the Diverse lineage and shows admixed ancestry (Fig. 3A-D). Increasing the number of *K* clusters in ADMIXTURE does not assign the single-leaf samples to a distinct ancestral group (Supplementary Figure S4). Instead, it highlights substructure within the five defined lineages. These results suggest that the seemingly new and distinct cluster observed in Figure 2 likely reflects admixture rather than a novel lineage.

**Figure 3.**
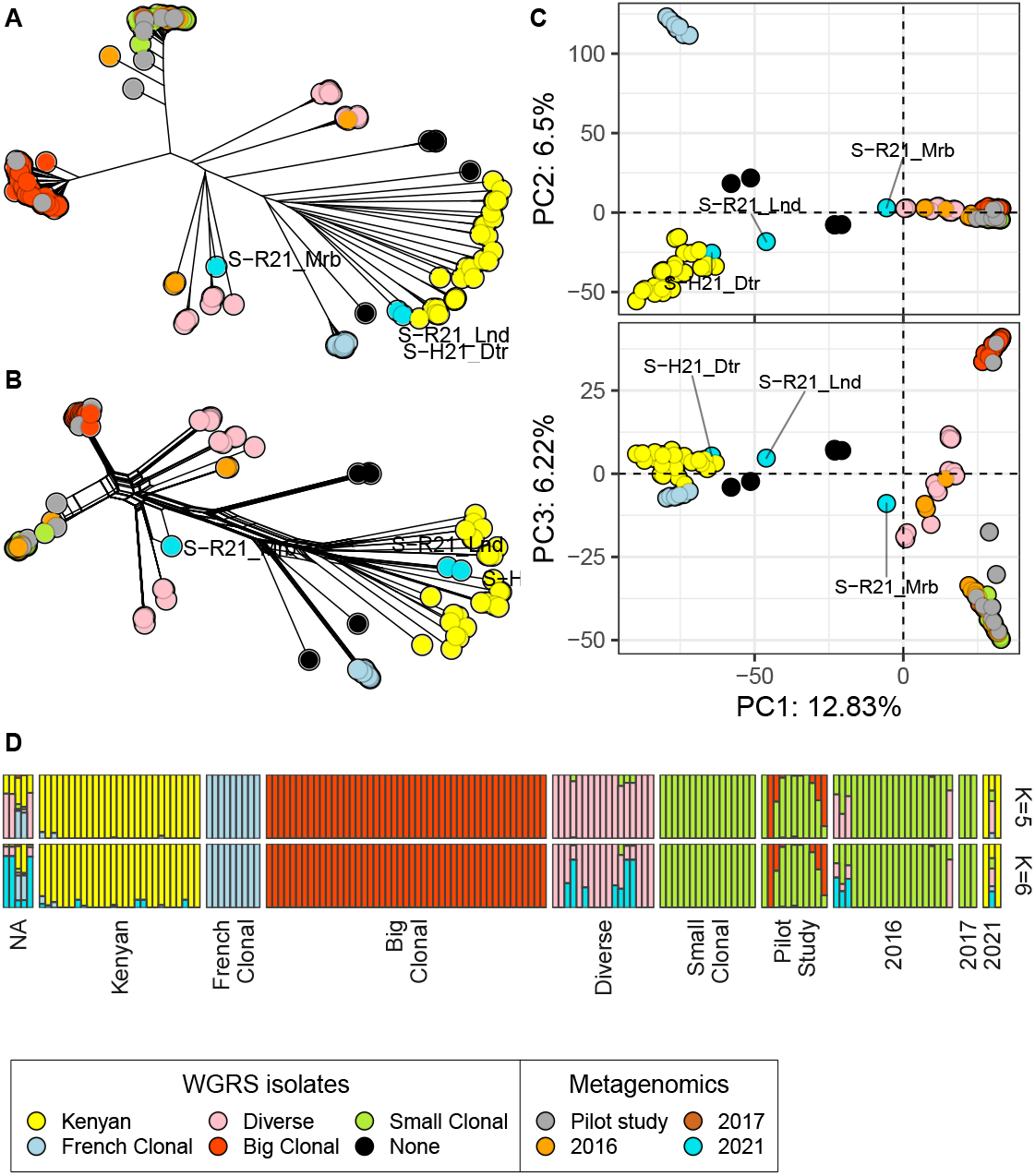
**A)** NJ tree, **B)** NNet **C)** PCA and **D)** ADMIXTURE ancestry coefficients of the metagenomic *E. turcicum* samples from single leaves with European and Kenyan isolates. Colors according to the five defined European and Kenyan WGRS lineages or the sampling year in the metagenomic samples.

### Relationship between extent of heterozygosity and mating type

To further assess whether the 2021 and 2022 samples represent admixed lineages, we calculated the percentage of heterozygous SNP calls per sample, with low values indicating infection by a single clonal lineage and higher values suggesting mixed-lineage infections (***Vidal-Villarejo et al., 2024***). Pooled samples from 2021 and 2022 show markedly higher heterozygosity than earlier metagenomic samples from 2016 and 2017, with the exception of two pooled samples (H21_Dtr and R22_Wdn) that exhibit low heterozygosity and only one mating type (Fig. 4). Among the three single-leaf samples from 2021, two show low heterozygosity and one shows elevated levels, yet all carry only one mating type. Our previous metagenomic analysis also identified a few mixed infections (in G7202_1, T7277_2 and R16_Bch3) with moderate heterozygosity, though still much lower than that observed in most pooled samples of the present study.

**Figure 4.**
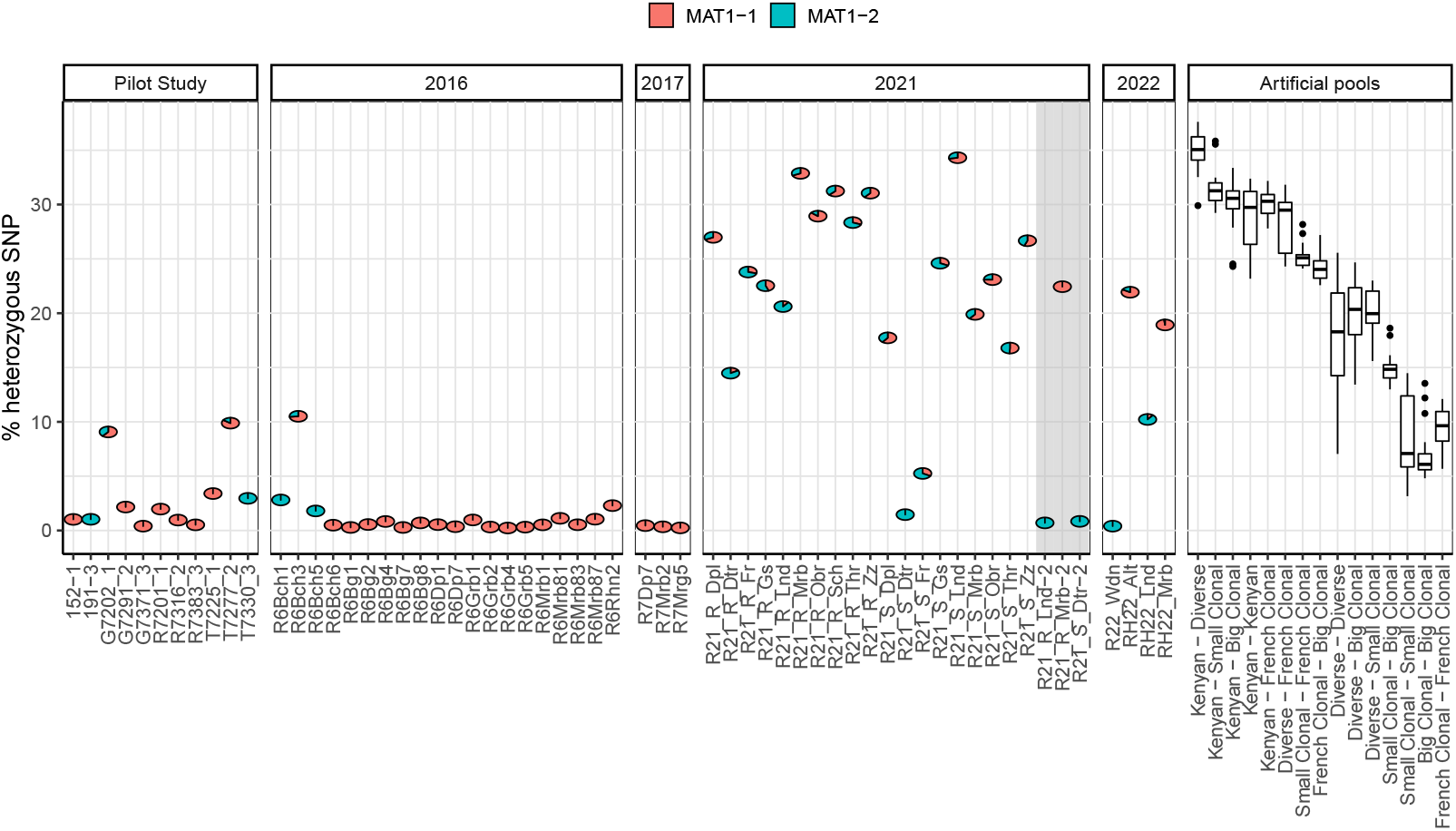
Percentage of heterozygous SNP. Pie charts indicate the percentage of reads mapping to each *MAT* mating type gene allele. Samples 152-1 and 191-3 are isolates (non-metagenomic) used for control. Single-leaf samples from 2021 are indicated with shaded background. Boxplots in “Artificial pools” correspond to the percentage of heterozygous SNP in artificial pools created with individuals from same and different clusters (2 individuals per cluster).

We hypothesize that the high heterozygosity in the pooled samples arises from the presence of *E. turcicum* strains belonging to genetically distant and diverse clusters, such as the Kenyan and Diverse lineages. Sequencing mixtures of such divergent and highly diverse lineages increases the proportion of heterozygous SNPs, particularly when the lineages share few common variants—for example, the Kenyan and European lineages share only 33% of SNPs (***Vidal-Villarejo et al., 2023***).

To test the impact of lineage mixing on heterozygosity, we created artificial pooled datasets by merging reads from WGRS isolates of the same or different *E. turcicum* lineages. These experiments confirmed that mixtures involving the Kenyan and Diverse lineages—especially those including multiple Kenyan isolates—consistently show the highest heterozygosity levels, closely matching those observed in metagenomic pooled samples (Fig. 4, Supplementary Fig. S5).

### Identification of different *E. turcicum* lineages

To further validate that the new, potentially admixed samples from 2021 and 2022 belong to one of the known lineages, we conducted additional analyses using all pooled and non-pooled metagenomic samples for comparison.

Using SNPmatch (***Pisupati et al., 2017***), we compared metagenomic samples to our reference panel of European and Kenyan WGRS isolates. SNPmatch showed 2021 samples and the four from 2022 had higher SNP match rates with isolates from the Kenyan, Diverse, or Small Clonal clusters, but none exceeded the statistical significance threshold for confident lineage assignment (Fig. 5A, Supplementary Fig. S6–S7). In contrast, the two 2019 samples and one from 2020 (H20_Sch) showed distinct match patterns consistent with the Small Clonal cluster, supporting their likely origin from this lineage, as also suggested by NJ tree analyses. To validate SNPmatch for metagenomic data, we applied it to the clonally infected, non-pooled samples from 2016 and 2017 and the pilot study, where it successfully assigned most samples to the Small Clonal or Diverse lineages consistent with their ADMIXTURE ancestry and low heterozygosity (Fig. 5A and Supplementary Figs. S8-S9). However, SNPmatch could not resolve the lineage composition of the three samples known to be admixed in 2016 and 2017 and the pilot study (Supplementary Figs. S8 and S9), indicating that while SNPmatch is effective at identifying lineages in samples infected with a single lineage, it struggles when samples contain multiple lineages.

**Figure 5.**
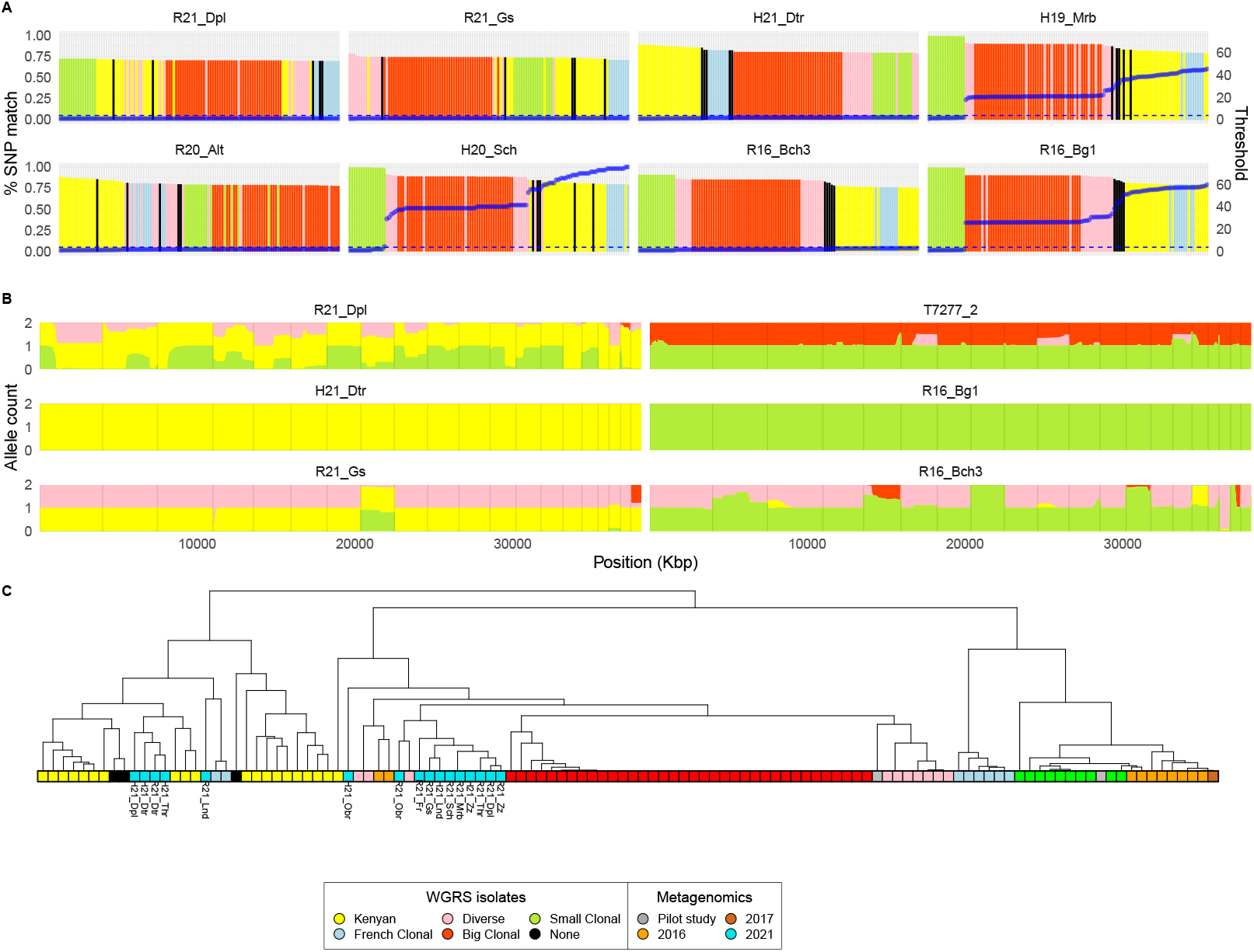
**A)** SNPmatch analysis. The left y-axis shows the percentage of SNPs matching the reference panel of WGRS isolates (x-axis) in eight metagenomic samples (full set in Supplementary Figs. S6–9). The right y-axis (blue) represents the likelihood ratio. The horizontal dashed blue line marks the 3.841 significance threshold. **B)** Local ancestry inference with ELAI in six metagenomic samples. ELAI plots indicate the source population for each of the two alleles along the genome (full set in Supplementary Fig. S10-12). **C)** Cladogram of metagenomic samples based on sequence coverage of protein-coding regions from genes differentiating Exserohilum lineages. Only samples with a minimum mean coverage of 10X are shown.

ADMIXTURE results for the three single-leaf samples from 2021 were skewed by the inclusion of pooled samples, prompting the use of local ancestry inference with ELAI (***Vi et al., 2023***) for more detailed analysis. Using the five genetic clusters of the European and Kenyan WGRS isolates as reference, ELAI revealed three main ancestry patterns among 2021–2022 samples (Fig. 5B and Supplementary Fig. S10): genomes dominated by Kenyan alleles; genomes split between Kenyan and Diverse clusters; and genomes admixed with alleles from Kenyan, Diverse, and Small Clonal lineages. In contrast, the non-pooled metagenomic samples of our previous study showed single cluster ancestry (mostly from Small Clonal) or known admixed profiles (Supplementary Fig. S10).

As a third approach to assign pooled samples to lineages, we performed hierarchical clustering based on the sequence coverage of previously identified presence-absence genes, including the mating-type locus *MAT1*, which differentiates *MAT1-1* and *MAT1-2* strains by presence or absence of sequencing coverage (***Vidal-Villarejo et al., 2023***). Using a 10X coverage threshold, 16 samples from 2021 were retained, five of which clustered with the Kenyan lineage (Fig. 5C) and were consistently classified as Kenyan by both ELAI (Fig. 5B, Supplementary Fig. S15) and ADMIXTURE at K=5 (Fig. 2B). Other samples clustered with a Diverse isolate, including one unclustered sample falling outside any main lineages (Fig. 5C). Lowering the threshold to 4X included three additional 2022 samples, one of which clustered with the Kenyan lineage and two with the Diverse group (Supplementary Fig. S14). In summary, all three analyses clearly indicated the presence of multiple pooled samples with an exotic, non-European ancestry.

### Spatiotemporal dynamics of *E. turcicum* across maize variety types

After characterizing the genetic lineages of infected leaf samples, we analyzed their spatiotemporal dynamics to evaluate changes in Kenyan ancestry across regions and variety types. Ancestry coefficients at *K* = 5 were averaged by location, with low-coverage samples imputed based on the five closest genetic neighbors. The *K* = 5 clustering level was considered sufficient given the absence of new lineages and consistency with ELAI results. By 2021, Kenyan ancestry exceeded 50% in nearly all samples (Fig. 6A) and locations (Fig. 6B), with mean values of 64% in Rheintaler Ribelmais and 73% in hybrid maize, while the Diverse lineage was more common in Ribelmais (mean 24%) than in hybrids (mean 12%). Small Clonal ancestry persisted in some fields, but the consistently high exotic Kenyan ancestry in 2022 indicates its establishment across the region.

**Figure 6.**
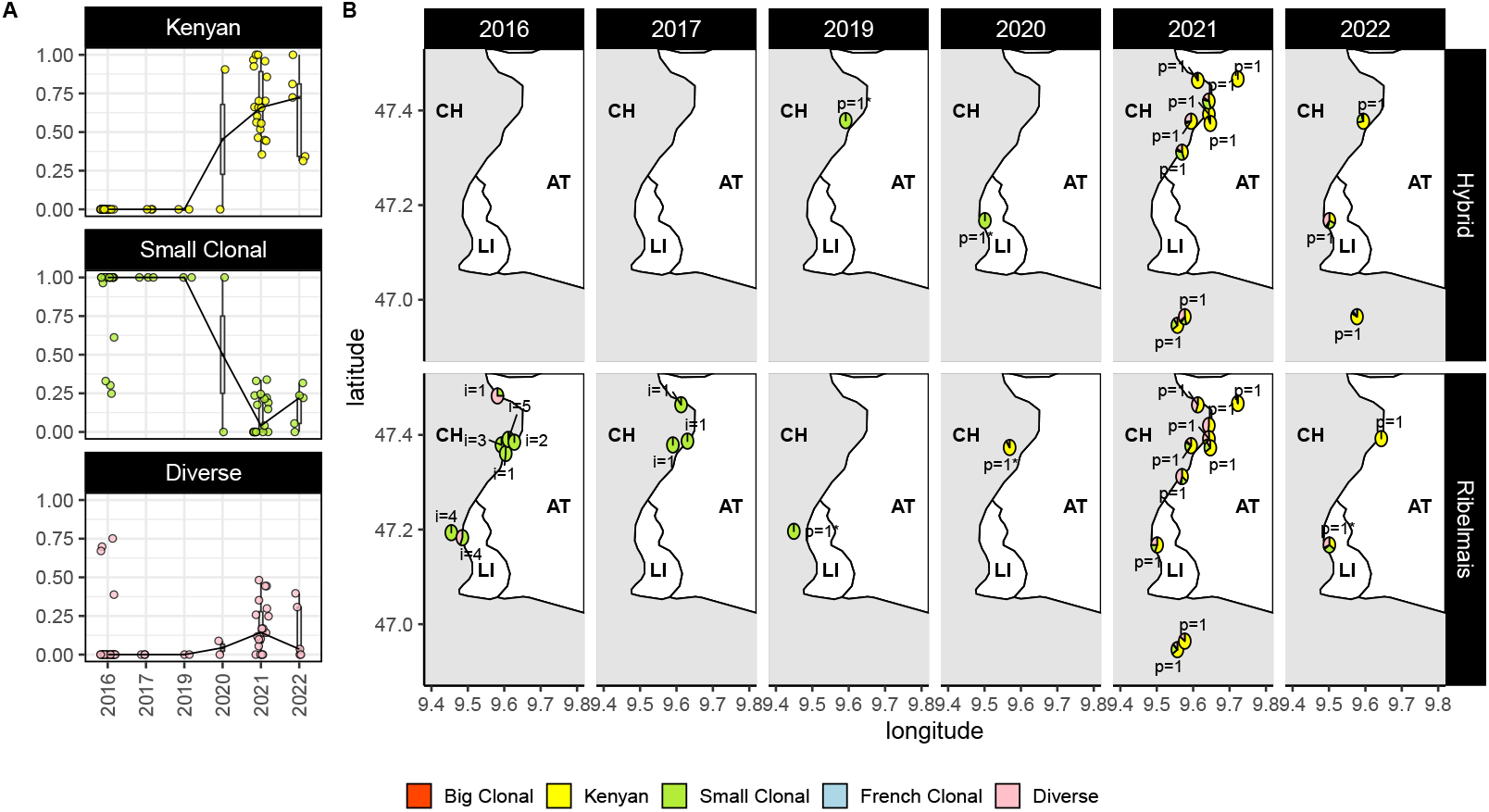
**A)** Boxplots (with overlaid jittered points) of ADMIXTURE ancestry coefficients at K = 5, across years, of the three most prevalent lineages in the region. The black line denotes the median ancestry proportion per year. **B)** Geographic distribution of ADMIXTURE ancestry coefficients at K = 5 summarized by location, year and maize variety. The text below pie charts indicate the number of single-leaf samples (i) or number of pooled samples (p). Asterisk denotes information built from imputed recovered samples.

### Differences in the phyllobiome

To assess differences in the phyllobiome composition of infected leaves across Rheintaler Ribelmais landrace, modern hybrid varieties, sampling year, and field locations, we conducted a multi-factorial Permutational Analysis of Variance (PERMANOVA) based on Bray-Curtis dissimilarities of taxonomically classified read abundances. The analysis revealed differences between years (***R*** = 0.45, *p* = 1*e*-05), between maize variety types with low explained variance (***R*** = 0.046, *p* = 0.002), and no variation among agricultural fields (*p* = 0.224; Table 2). A pairwise PERMANOVA showed that the greatest interannual deviation occurred in 2021 (***R*** = 0.40 ~ 0.50, *p* = 0.001; Table 2). It can also be observed in the Principal Coordinates Analysis (PCoA), where the first axis clearly distinguishes most 2021 samples and three 2022 samples from the other years (Fig. 7A), which correspond to the samples with highest *Exserohilum* percentage in Fig. 1. Due to the pronounced differences in 2021, we conducted year-specific PERMANOVA analyses, which confirmed that phyllobiome differences between maize variety types were not statistically significant within any year (*p* > 0.01 in all cases; Supplementary Fig. S15).

**Table 1.** Summary of 60 pooled samples over year and variety type. Each samples includes cuttings of infected leaves from at least randomly selected plants from a single agricultural production field.

| Sampling year | Collection sites |  |  | Retained samples |  |  |
| --- | --- | --- | --- | --- | --- | --- |
|  | Rheintaler | Hybrids | Total | Rheintaler | Hybrids | Total |
| 2019 | 9 | 5 | 14 | 1 | 1 | 2 |
| 2020 | 8 | 4 | 12 | 1 | 1 | 2 |
| 2021 | 10 | 10 | 20 | 10 | 9 | 19 |
| 2022 | 7 | 7 | 14 | 2 | 3 | 5 |

**Table 2.** Multifactorial PERMANOVA test statistics for significance of factors influencing phyllobiome dissimilarities and pairwise PERMANOVA on the year.

| Source of variation | df | SS | MS | pseudo-F | *R*<sup>2</sup> | *p* value |
| --- | --- | --- | --- | --- | --- | --- |
| Year | 3 | 1.083 | 0.361 | 17.87 | **0.452** | 1e-05 |
| Variety type | 1 | 0.110 | 0.110 | 5.432 | **0.046** | 0.002 |
| Site | 13 | 0.304 | 0.023 | 1.169 | 0.127 | 0.224 |
| Residuals | 45 | 0.900 | 0.020 | - | 0.376 | - |
| Pairwise groups by year |  |  |  |  |  |  |
| 2019, 2020 | 1 | 0.090 | 0.090 | 3.218 | **0.118** | 0.003 |
| 2019, 2021 | 1 | 0.681 | 0.681 | 35.627 | **0.504** | 0.001 |
| 2019, 2022 | 1 | 0.080 | 0.080 | 3.217 | **0.110** | 0.002 |
| 2020, 2021 | 1 | 0.567 | 0.567 | 28.078 | **0.460** | 0.001 |
| 2020, 2022 | 1 | 0.058 | 0.058 | 2.158 | 0.082 | 0.039 |
| 2021, 2022 | 1 | 0.462 | 0.462 | 25.242 | **0.419** | 0.001 |

df: degrees of freedom; SS: Sum of Squares; MS: Mean Squares The analysis was performed with 99,999 permutations and calculated on Bray-Curtis dissimilarities. Values indicated in bold are *R*<sup>2</sup> with *p* value ≤ 0.01. Site refers to collection site.

**Figure 7.**
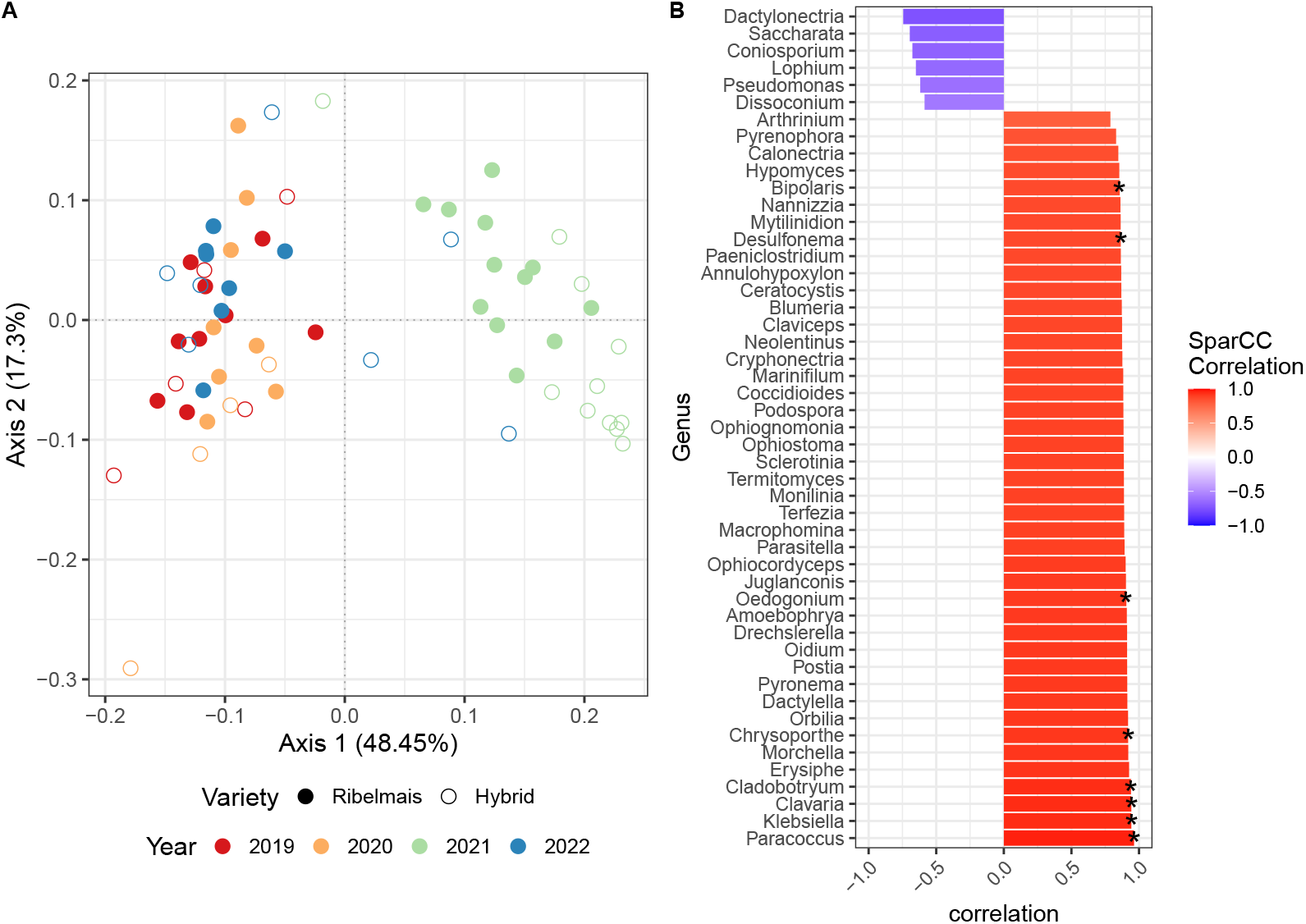
Analysis of phyllobiome composition based on metagenomic sequencing data. **A)** Principal Coordinate Analysis of the Bray-Curtis dissimilarities of phyllobiome composition. **B)** SparCC correlation coefficients of taxa with direct interaction with *Exserohilum*. The asterisk (*) in B denotes the interaction detected in both the Neighborhood and Inverse Covariance methods of SPIEC-EASI.

### Correlation of other microbial taxa with *E. turcicum*

To identify positive and negative associations between microbial taxa and *Exserohilum*, such as the previously observed relationship with *Klebsiella* (Fig. 1), we applied SPIEC-EASI (***Kurtz et al., 2015***). This method is designed for compositional data and infers direct associations while minimizing indirect effects. Since SPIEC-EASI edge weights are not directly interpretable as correlation coefficients, we further quantified the strength of these associations using SparCC (***Friedman and Alm***, ***2012***), another method tailored for compositional microbiome data. Focusing on *Exserohilum*-related interactions, SPIEC-EASI identified 43 taxa with positive associations, including *Klebsiella* as shown in Fig. 1. In addition, six taxa were found to have direct negative associations with *Exserohilum* (Fig. 7B), including *Pseudomonas, Lophium, Coniosporium, Saccharata*, and *Dactylonectria*.

## Discussion

Our study builds on earlier monitoring of *E. turcicum* diversity in the Swiss Rhine Valley conducted between 2016 and 2018 (***Vidal-Villarejo et al., 2024***), extending it by incorporating samples from additional years and implementing a new pooling strategy to reduce sequencing costs. Infected maize leaf samples were collected annually from 2019 to 2022 following visual inspection for signs of *E. turcicum* infection. Unlike in the previous study, several samples in the current survey that lacked *Exserohilum* reads showed high abundance of *Puccinia* instead, the causal agent of common rust in maize; these samples were excluded from further analysis of *Exserohilum* diversity.

### A pooling strategy with an improved bioinformatics pipeline

We improved the bioinformatics pipeline for monitoring *Exserohilum* lineages compared to our previous survey (***Vidal-Villarejo et al., 2023***). Instead of applying the Kaiju metagenomic classifier to all sequencing reads, we first filtered out those mapping to the maize genome, which reduces computational load and processing time. For lineage detection, we retained core methods such as NJ tree, NNet, PCA, ADMIXTURE, heterozygosity, and mating type analyses. To improve classification robustness, we additionally integrated three new approaches: SNPmatch (***Pisupati et al., 2017***), ELAI (***Vi et al., 2023***), and hierarchical clustering based on protein-coding presence-absence genes. By pooling ten infected leaves per field into a single metagenomic sample, we reduced both sequencing and laboratory costs by a factor of ten, while maintaining comparable resolution to single-leaf data. We also developed a recovery approach for samples with very low proportions of *Exserohilum* reads, which would otherwise be excluded under standard filtering thresholds.

The pooling strategy did not fully resolve the issue of low pathogen read counts in a substantial number of samples. A likely explanation is that lesions caused by *E. turcicum* were mistakenly sampled alongside those from other pathogens, as indicated by samples dominated by *Puccinia* or *Alternaria* reads. To increase the accuracy of future sampling, either additional training of field personnel or automated lesion recognition using image analysis (e.g., ***Ahmad et al., 2023***) may improve identification of *E. turcicum* infections. Future strategies should aim to streamline sample preparation while ensuring sufficient amounts of *Exserohilum* DNA in samples for reliable, cost-effective analysis—especially given the logistical challenges of processing hundreds of field samples under time constraints.

### Identification of a new genetic lineage

The presence of multi-lineage infections with an exotic genetic ancestry of the Kenyan cluster led to the appearance of a novel genetic group in population structure analyses of the pooled samples, potentially causing misclassification. For example, PCA results conflicted with NJ tree and ADMIXTURE analyses, as samples scattered between the Diverse and Kenyan clusters rather than forming a distinct group. Pooled samples also showed unusually high heterozygosity compared to single-leaf metagenomic samples—a pattern inconsistent with previously observed single- or multi-lineage infections and suggestive of sequencing artifacts rather than a new clonal lineage. To validate this, we sequenced three individual leaves from pooled samples; two showed reduced heterozygosity, while one remained highly heterozygous, indicating a true mixed infection involving genetically divergent lineages such as Kenyan and Diverse.

These findings confirmed the mixed, non-clonal nature of the pooled samples. SNPmatch successfully distinguished single-from multi-lineage infections, while ELAI identified genomic mixtures involving Kenyan, Diverse, and/or Small Clonal ancestry. A third method—classification based on lineage-specific presence-absence genes—was less robust due to its reliance on high sequencing coverage. Given the limitations of metagenomic data, the latter approach may introduce errors and could be excluded from future workflows.

### Decline of clonal lineages in favor of lineages with Kenyan ancestry

Despite the limited number of samples with sufficient *E. turcicum* reads, we obtained enough data to investigate the temporal dynamics of genetic lineages in the Swiss Rhine Valley. All samples from 2021 and 2022 belonged to the Kenyan lineage or showed admixture with the Kenyan, Diverse, or Small Clonal clusters. Whereas the Small Clonal lineage dominated the region in 2016 and 2017 (***Vidal-Villarejo et al., 2024***), it is no longer predominant, having been largely replaced by the lineages from the exotic Kenyan cluster. Although low read counts and few samples from 2018–2020 create resulted in missing data from these years, available evidence suggests that the transition from Small Clonal to Kenyan and Diverse lineage mixtures occurred between 2019 and 2020, with samples from 2022 confirming continued Kenyan dominance. Our initial classification of five clonal lineages in Europe (***Vidal-Villarejo et al., 2023***) was based on collections from 2012–2014, during which the Kenyan cluster was mostly limited to samples from Kenya and a single isolate from Turkey. Five additional European isolates showed ancestry with this exotic cluster, but their admixture with Diverse or French Clonal alleles precluded their assignment to a specific lineage (see ‘NA’ group in Fig. 2C and Supplementary Figures S2 and S4). These admixed isolates differ from the metagenomic samples from 2021 and 2022 in their ADMIXTURE ancestry coefficients (Fig. 2C, Supplementary Figures S2 and S4), but the presence of Kenyan ancestry in current samples may reflect either the local spread of previously admixed isolates or new introductions from outside Europe. A continent-wide screening of current *E. turcicum* populations would help determine whether the rise of the exotic Kenyan lineage is widespread or localized to the Rhine Valley. Ongoing monitoring is crucial to assess whether this lineage will become established or be replaced by other existing or emerging lineages. Given the rapid shifts observed in *E. turcicum* population structure, our findings highlight the importance of sustained surveillance to detect emerging threats.

### Differences in pathogen diversity between variety types

The multiyear monitoring of *E. turcicum* has revealed a shift from the Small Clonal lineage to the Kenyan lineage (Fig. 6A). To determine whether this shift occurred similarly in both the highly susceptible Rheintaler cultivar and hybrid varieties bred for NCLB resistance, we analyzed the admixture coefficients separately for each variety type. Our analysis showed comparable proportions of the Kenyan lineage in both, although the Diverse lineage appeared more pronounced in the Rheintaler landrace than in the hybrid varieties (Fig. 6B). While it remains to be seen whether these differences will become stronger over time, our findings suggest that the susceptible Rheintaler Ribelmais landrace could serve as a useful ‘sentinel’ genotype for the early detection of new pathogen strains (***Lovell-Read et al., 2023***).

### Metagenomic reads highlight taxa of interest associated with *E. turcicum*

Analysis of phyllobiome composition revealed no significant differences between the two maize varieties, despite previous studies reporting host genetic effects on leaf microbiomes in maize (***Wallace et al., 2018; Wagner et al., 2020***) and other crops (***VanWallendael et al., 2022; Morris and Bohannan, 2024***). This lack of differentiation may be attributed to our targeted sampling strategy. Notably, phyllobiome composition in 2021 differed significantly from other years and coincided with the highest number of *Exserohilum* reads. Unlike earlier studies where interannual variation was linked to environmental differences or sampling time, all samples in this study, except two collected in July 2019, were collected in mid-September, making sampling date an unlikely explanation. Furthermore, phyllobiome composition did not significantly vary across collection sites.

The metagenomic survey also enabled an association analysis with *Exserohilum*. Using SPIEC-EASI, we identified six taxa negatively associated with *E. turcicum*, including the genus *Pseudomonas*, which is widely used as a biocontrol agent (BCA) due to its antifungal properties (***Haas and Défago, 2005; Weller, 2007; Antolín-Llovera et al., 2012***). Consistent with this, *Pseudomonas fluorescens* was previously shown to inhibit *E. turcicum* in vitro (***Mostafa Hassan, 2017***). A strong negative correlation with the yeast *Metschnikowia* was observed in our earlier study (***Vidal-Villarejo et al., 2024***), but not in the recent data from 2020 and 2021 because *Metschnikowia* did not pass tresholds of minimum relative DNA abundance. SPIEC-EASI also identified several taxa positively associated with *Exserohilum*, including *Klebsiella*; these associations may reflect shared environmental conditions or potential microbial facilitation.

These findings highlight taxa of potential interest due to their contrasting associations with *E. turcicum*. The variability in microbial correlations suggests that controlled experiments—such as those using synthetic communities (SynComs) (***Xu et al., 2025; Durán et al., 2025***)—are needed to determine whether these associations represent causal interactions between the pathogen, its microbiome, and the host genotype. Such insights could support breeding strategies or the development of microbiome-based plant protection measures (***Hawkes et al., 2021***).

### Perspectives for the monitoring of *E. turcicum*

High-throughput sequencing (HTS) of pooled samples enables the efficient survey of large numbers of individual plants, enhancing the potential of metagenomic approaches for monitoring pathogen dynamics. A major advantage of metagenomic HTS over traditional methods is its ability to detect novel strains while generating sufficient data to study pathogen demography, evolution, and host interactions (reviewed by ***Nizamani et al., 2023***). Achieving this goal requires a well-curated global reference dataset of fully sequenced genomes, which already exists for fungal pathogens such as *Magnaporthe oryzae* (***Gladieux et al., 2018; Ali et al., 2023***), *Ramularia collo-cygni* (***Stam et al., 2019***), *Pyrenophora teres* f. *teres* (***Taliadoros et al., 2024***), and *Zymoseptoria tritici* (***Feurtey et al., 2023***). For *E. turcicum*, such a reference dataset is currently lacking, limiting our ability to precisely infer the geographic origin of samples clustering with the exotic Kenyan lineage. Although these samples likely belong to a genetically diverse tropical group, their exact origin remains uncertain and could also be Central or South America, where the pathogen is believed to have evolved (***Borchardt et al., 1998b***).

This information would be highly valuable for breeding resistance against Northern Corn Leaf Blight (NCLB), as tropical pathogen races may overcome resistance genes used in temperate breeding programs. Breeding strategies focused on quantitative resistance could offer more durable protection against both temperate and tropical races (***Schechert et al., 1999; Welz and Geiger, 2000; Galiano-Carneiro et al., 2021***). Metagenomic monitoring could help identify emerging pathogen races adapted to a changing climate, thereby informing resistance breeding. In addition, bioinformatic analyses may aid in the discovery and characterization of novel virulence genes, supporting evolutionary and functional studies to model the epidemiology of *E. turcicum*.

## Material and Methods

### Leaf sampling and sequencing

Leaf samples were collected between 2019 and 2022 by M. H. and field technicians. In mid-September, maize fields in the Rhine Valley, spanning from Lake Constance in the north to Landquart in the south, were sampled. From each field, one *Exserohilum*-infected leaf was collected from 25 individual plants distributed diagonally, excluding the six rows sown transversely at the field edge. The nearest hybrid maize field was sampled using the same scheme, with edge rows included in case too few infected plants were found in the inner rows of the field. Lesions suspected to be caused by *E. turcicum* were excised with scissors from the collected leaves and dried on filter paper with silica gel. Samples were labeled with the collection date and GPS coordinates. Sample names follow a structured format: (i) maize variety, denoted as ‘R’ for Ribelmais and ‘H’ for Hybrid, (ii) year of sampling, represented by the last two digits of the year (e.g., ‘16’ for 2016), and (iii) collection site, abbreviated to three letters. DNA was extracted from 10 randomly selected leaves per field out of the 25 collected and prepared for metagenomic short-read sequencing using the same protocol as in (***Vidal-Villarejo et al., 2024***). Samples were sequenced in three separate batches using Illumina NovaSeq. The sequencing was performed in three batches. The first batch included 20 pooled samples from 2021 and generated a total of 205.5 Gbp of sequencing data. The second batch consisted of three single-leaf (non-pooled) samples from 2021, yielding 46 Gbp. The third batch comprised 40 pooled samples from 2019, 2020, and 2022, producing a total of 424.8 Gbp of sequencing data.

### Metagenomic sequences processing and variant calling of *E. turcicum* reads

A Snakemake pipeline was developed to process metagenomic sequencing reads and identify nucleotide variants in *E. turcicum* specifically for the pooled samples. Initially, the pipeline used TrimGalore (v0.6.7; ***Krueger, 2015***) for read trimming and quality filtering but was later modified to use fastp (v0.22.0; ***Chen et al., 2018***). Samples from the first sequencing batch were processed with TrimGalore, while the rest were handled with fastp.

After read cleaning, the pipeline maps the reads to the maize B73 reference genome V4 (AGPv4; ***Jiao et al., 2017***) using BWA (v0.7.17; ***Li and Durbin, 2009***). Reads that do not align with the host genome are taxonomically classified with Kaiju (v1.7.4; ***Menzel et al., 2016***) using the pre-built database ‘nr_euk’ (v2022-03-10; provided by the Kaiju authors), which includes the non-redundand database containing archea, bacteria, viruses, fungi and microbial eukaryotes. Genus-level Kaiju classifications are obtained using the *kaiju2table* script included in Kaiju. Reads classified as *E. turcicum* (taxonomic IDs 671987 and 93612) are then mapped to the reference genome *Setosphaeria turcica* Et28A v1.0 (***Ohm et al., 2012; Condon et al., 2013***) and to the two MAT sequences (GenBank IDs GU997138.1 and GU997137.1) to determine their mating type. Reads aligning to the *E. turcicum* genome are realigned with GATK (v3.7.0; ***McKenna et al., 2010***), and duplicates are removed using the Picard tool suite (v2.18.23, http://broadinstitute.github.io/picard/). Reads containing more than five mismatches are filtered out with BAMtools (v2.5.1; ***Barnett et al., 2011***), as they may result from incorrect Kaiju classifications. Finally, single nucleotide variants were identified with BCFtools (v1.11; ***Narasimhan et al., 2016***) using a predefined list of variant positions corresponding to those in the VCF file with data from (***Vidal-Villarejo et al., 2024***). This restriction ensures that variants detected in metagenomic samples are true *Exserohilum* variants rather than sequences from taxonomically similar species that may also align to the *Exserohilum* genome.

The resulting VCF file was merged with a VCF file with data from (***Vidal-Villarejo et al., 2024***), which contains 10,333 SNP from 121 *E. turcicum* isolates from Europe and Africa, along with 41 metagenomic *E. turcicum* samples from the Rhine Valley. This merged VCF file was filtered first by excluding SNPs with more than 40% missing data and those with more than two alleles, resulting in a VCF set of 10,260 positions and 225 samples. With the resulting VCF file, a second additional filtering was applied: A minimum of two reads was required for a genotype to be called, loci with more than 20% missing data were removed, and samples with more than 50% missing loci were excluded.

### Filtering of samples with low coverage of *E. turcicum*

We designed an alternative filtering strategy to accommodate low-coverage samples. Starting with the VCF file containing 10,260 SNPs and 225 samples, we retained loci with no missing data in the low-coverage samples, to ensure that the selected SNP set was informative for the recovered samples. To further reduce missing data, we then removed loci with more than 20% missing data and excluded samples with more than 20% missing loci. This strategy was applied separately for each of the five recovered samples. Each recovered sample resulted in a new separated VCF file.

### Identification of mating type and genetic ancestry

To classify and assess the distribution of MAT types, we counted the reads mapped to the *MAT1-1* and *MAT1-2* sequences. We then calculated the proportion of each MAT type by dividing the number of reads mapped to a specific allele (*MAT1-1* or *MAT1-2*) by the total number of reads mapped to either sequence. To confidently assign a MAT type to a sample, we required at least 50% sequence coverage by one or more reads. Based on this criterion, samples were classified as carrying one or both MAT types, with the calculated percentage reported for each.

A NJ tree (***Saitou and Nei, 1987***) was constructed from Euclidean distances using allele frequencies derived from read counts, employing the R package ‘ape’ (v5.8; ***Paradis and Schliep, 2019***). The number of heterozygous SNPs per sample was calculated by dividing the total number of heterozygous SNPs by the number of non-missing positions. Artificial pooled sequences were generated by merging WGRS sequencing read counts from randomly selected isolate samples within and across the five genetic clusters. Twenty artifial pools of random samples were generated for each pairwise group of clusters. We created artifical pools with various number of samples per cluster, from 1 sample per cluster to 6. For these artificial pools, the percentage of heterozygous SNPs was calculated using the same method as for the metagenomic samples. Population structure inference with ADMIXTURE was performed using 20 independent runs with *K* = 1 to 10 clusters. The results were merged with CLUMPACK, and the ‘Major Cluster’ (the most common clustering arrangement) was used as the summarized outcome of the different runs. A NeighborNet was constructed from pairwise Nei’s distances applying a pairwise exclusion approach. Nei’s distances were imported into SplitsTree6 to generate the NeighborNet (***Bryant and Moulton, 2004; Huson and Bryant, 2006***). A Principal Components Analysis (PCA) was calculated with the R package ‘ade4’ (***Dray and Dufour, 2007***) and using allele frequencies derived from read counts, with missing loci imputed to the mean genotype, as PCA does not allow for missing data.

To identify different *E. turcicum* strains, we used SNPmatch, a tool that compares query genotypes against a reference panel to determine the most likely match based on SNP similarity and likelihood ratios (***Pisupati et al., 2017***). The reference panel, constructed using the *makedb* option, included WGRS data from European and Kenyan isolates, with haploid genotypes converted to homozygous diploids to meet the requirement of diploid sequences as input. Since SNPmatch processes each query sample separately, we opted to use the merged SNP dataset of 10,260 variants instead and apply filtering criteria separately for each sample, requiring a minimum read depth of 2 and 0% missing data. This approach aimed to retain a higher number of SNPs per sample, as each sample had to be analyzed separately. SNPmatch was run individually for each sample using the *inbred* option, which calculates the percentage of matching SNPs and a likelihood ratio against the best hit. A likelihood ratio below 3.841—assuming a chi-squared distribution—indicates no significant difference between the top matching reference genotype and the rest of reference samples.

Local admixture was estimated using ELAI (***Vi et al., 2023***) with the first 20 scaffolds of the reference assembly. European and Kenyan clusters were used as source populations. For each source population, a VCF file was extracted using VCFtools (v0.1.16, ***Danecek et al., 2011***) and converted into BIMBAM format with PLINK (v1.9, ***Purcell et al., 2007***). ELAI was run 10 times independently, each time using five different generation times (5, 10, 15, and 20) with the following options: `-s 30 -C 5 -c 25 -mg [5,10,15,20] –exclude-nopos –exclude-maf 0.01 –max-missing1`. To summarize the results, we averaged across all 10 independent runs and the five generation times into a single final estimate.

To cluster samples based on presence-absence genes, we first calculated the sequence coverage of *E. turcicum* protein-coding sequences (CDS), as Kaiju classifies only reads from protein-coding regions. We then extracted sequence coverage for a set of 62 *E. turcicum* genes previously identified in ***Vidal-Villarejo et al***. (***2023***) as presence-absence genes distinguishing the European and Kenyan lineages. A position was considered covered if it had at least one read. To prevent clustering due to coverage differences, we applied a minimum mean coverage threshold of 4X and 10X per sample. Samples were clustered based on gene coverage percentage using the R packages ‘pheatmap’ (***Kolde, 2019***) and ‘stats’ (***R Core Team, 2018***).

### Phyllobiome composition and correlations with *Exserohilum*

To analyze phyllobiome composition, we first filtered the taxonomic classifications obtained with Kaiju at the genus level by excluding taxa with < 0.01% relative abundance in fewer than five samples and samples with < 1, 000 reads. A multi-factorial permutational analysis of variance (PERMANOVA) was conducted for three factors (year, maize variety, and collection site) using the Bray-Curtis distance of the square root-transformed relative abundances of Bacteria and Eukarya using the following model

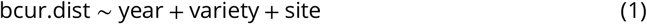

with 99,999 permutations, using the R package ‘vegan’ (***Oksanen et al., 2019***). Pairwise PERMANOVA for year effects was performed with the R package ‘pairwiseAdonis’ (***Martinez Arbizu, 2017***). Group dispersion homogeneity was tested using the *betadisper* function from the R package ‘vegan’ with 99,999 permutations.

Direct associations among taxa and *Exserohilum* were identified using the R package ‘SPIECEASI’ (***Kurtz et al., 2015***). This package includes a function to also calculate SparCC correlation from ***Friedman and Alm (2012)***. The Neigbourhood selection (mb) and inverse covariate (glasso) methods from SPIEC-EASI were both calculted with 40 lambda, 50 repetitions, and a minimum of 0.01 to identify the direct relationships between the microbiome and *Exserohilum*. Thereafter, direct relationships were weighted with the correlation coefficients from SparCC.

## Author Contributions

M.V-V. and K.S designed the study. B.K, M.H., B.O and H.O designed leaf sampling strategy, collected and provided leaf material. M.V-V conducted the data analyses. M.V-V and K.S. interpreted the data and wrote the manuscript. All authors read and agreed to the final version of the manuscript.

## Acknowledgments

We are grateful to Vanessa Haseneder for DNA extraction and sequencing library construction. Leaf sampling was aided by members of the Verein Rheintaler Ribelmais and employees of RhyTOP GmbH. This work was supported by the Bundesministerium für Forschung, Technologie und Raumfahrt (BMFTR) under grant number 031B0731A as part of the research program ‘Agricultural systems of the future’, project ‘NOcsPS - A more Sustainable Agriculture 4.0 without chemical-synthetic Plant Protection’. The authors acknowledge the additional support by the state of Baden-Württemberg through bwHPC. The authors declare no conflict of interest.

## Data Availability

The data that support the findings of this study are openly available. Raw sequencing data were deposited to the European Nucleotide Archive (ENA; https://www.ebi.ac.uk/ena) under project number PRJEB90581. Derived sequencing data like VCF files were deposited and scripts were submitted to Zenodo doi:10.5281/zenodo.15711400 [Accessible after publication].

