## Supporting Information S1 for "Rapid replacement of established by exotic genetic lineages of the fungal maize pathogen *Exserohilum turcicum* in the Swiss Rhine valley"

### Replacement of *Exserohilum turcicum* clonal lineages in the Rheintaler Swiss Valley by highly diverse populations from Europe and Africa

**Mireia Vidal-Villarejo**[1], **Benedikt Kogler**[1], **Michael Hammerschmidt**[2], **Barbara Oppliger**[2], **Hans Oppliger**[2], **Karl Schmid** [1] ✉

[1]Institute of Plant Breeding, Seed Science and Population Genetics, University of Hohenheim, Stuttgart, Germany; [2]Landwirtschaftliches Zentrum Sankt Gallen, Salez, Switzerland; [3]Verein Rheintaler Ribelmais e.V.

✉ **For correspondence:**
 (KJS)

#### – Supporting Information –

#### Contents

**Supplementary Figures** **2**

### Supplementary Figures

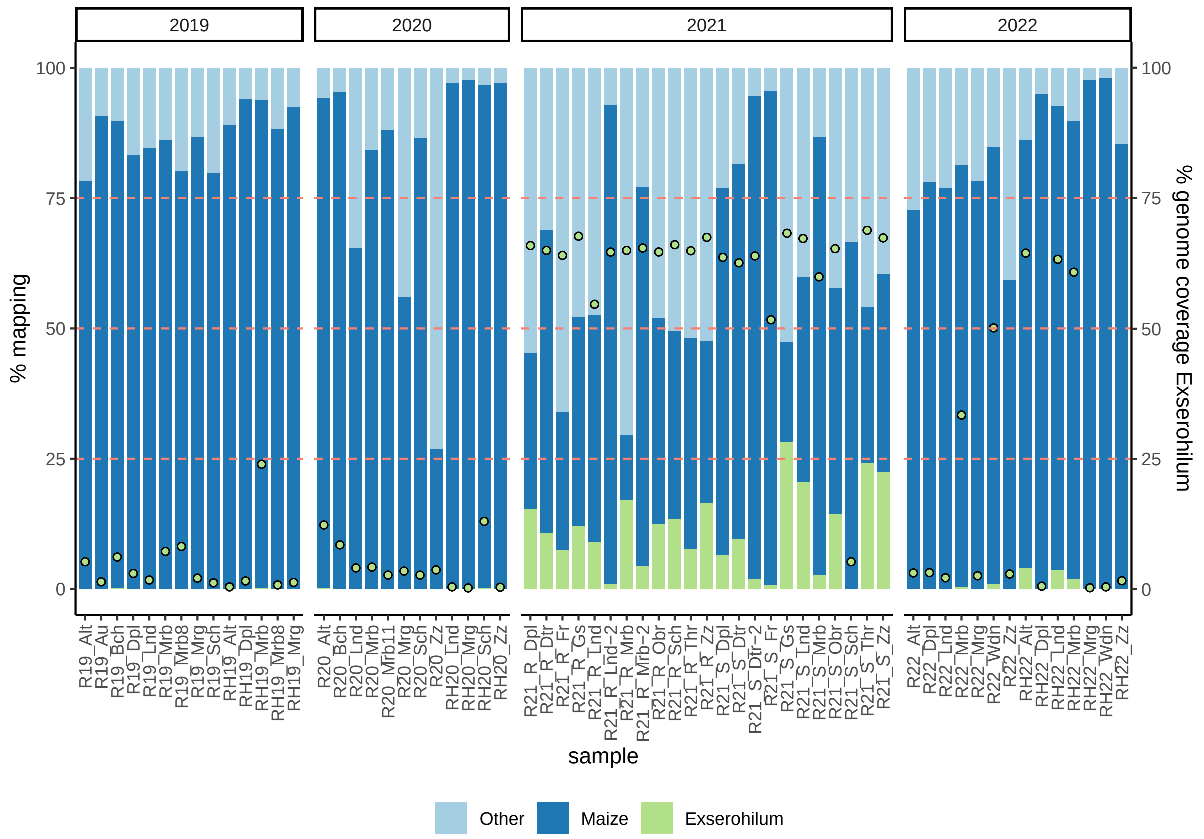


**Figure S1.** Percentage of reads mapping (left y-axis) to maize and Exesrohilum. "Other" represents reads not mapping to any of the two genomes. Dots (right y-axis) indicate the percentage of genome covered by at least one read in Exserohilum genome

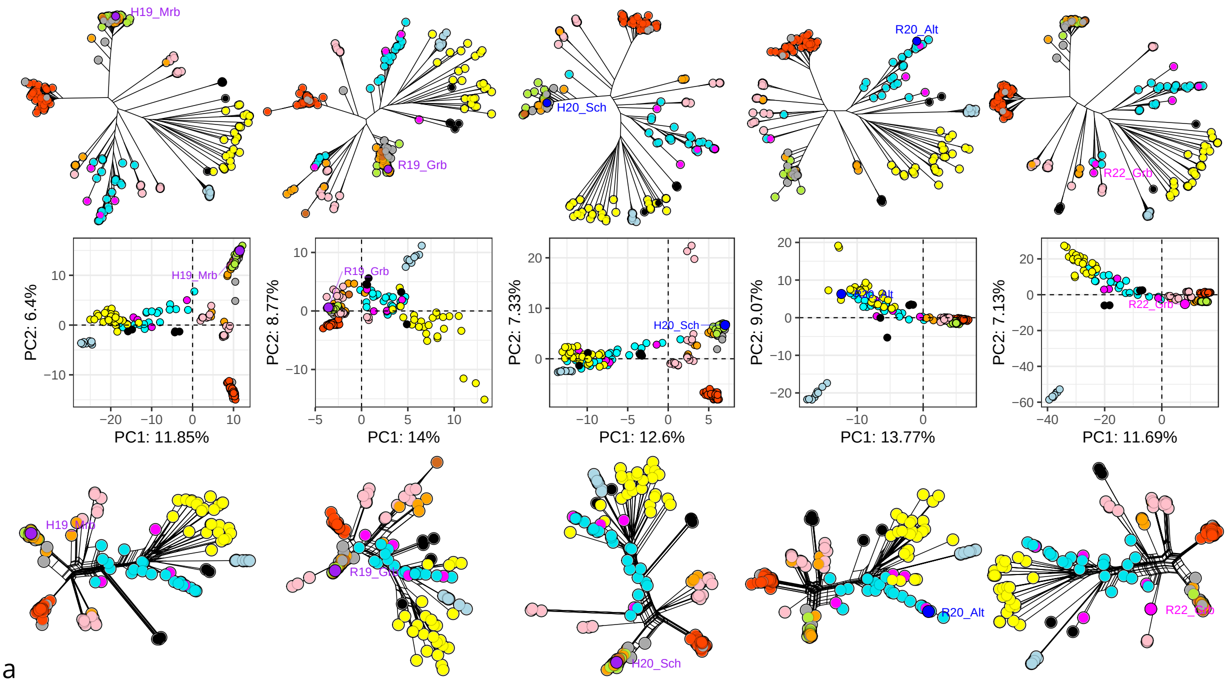


**Figure S2.** NJ Tree (top), PCA (middle) and NNet (bottom) of the five recovered samples, indicated by their name

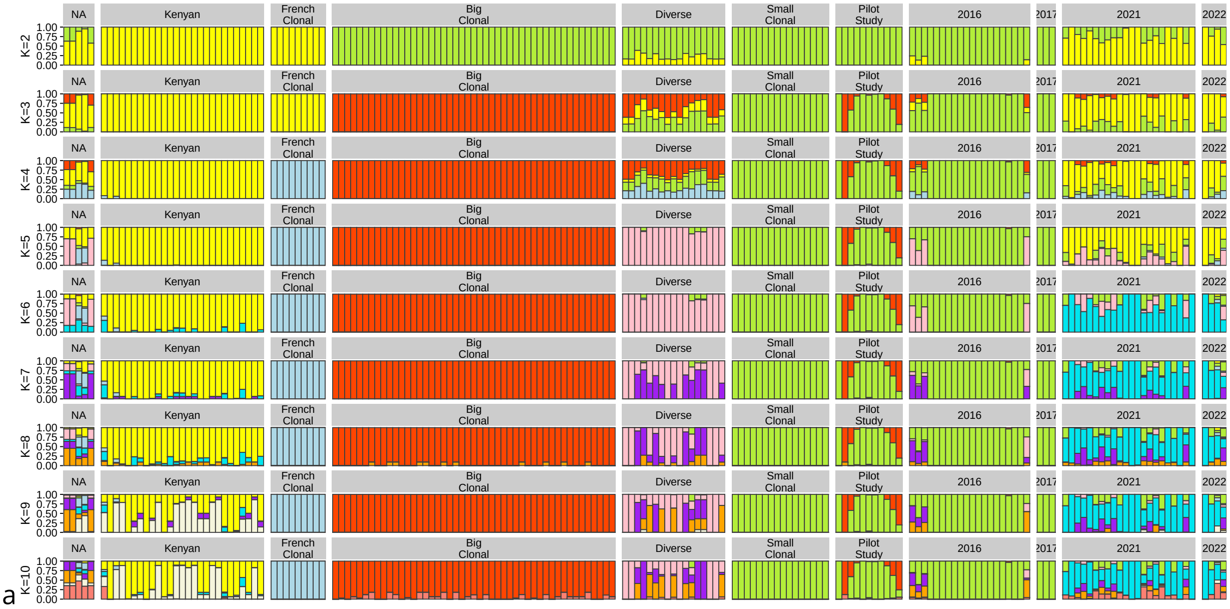


**Figure S3.** Admixture ancestry coefficients in K from 2 to 10, including metagenmic samples (indicated by the year of sampling or pilot study) and WGS Isolates (indicated by their lineage name)

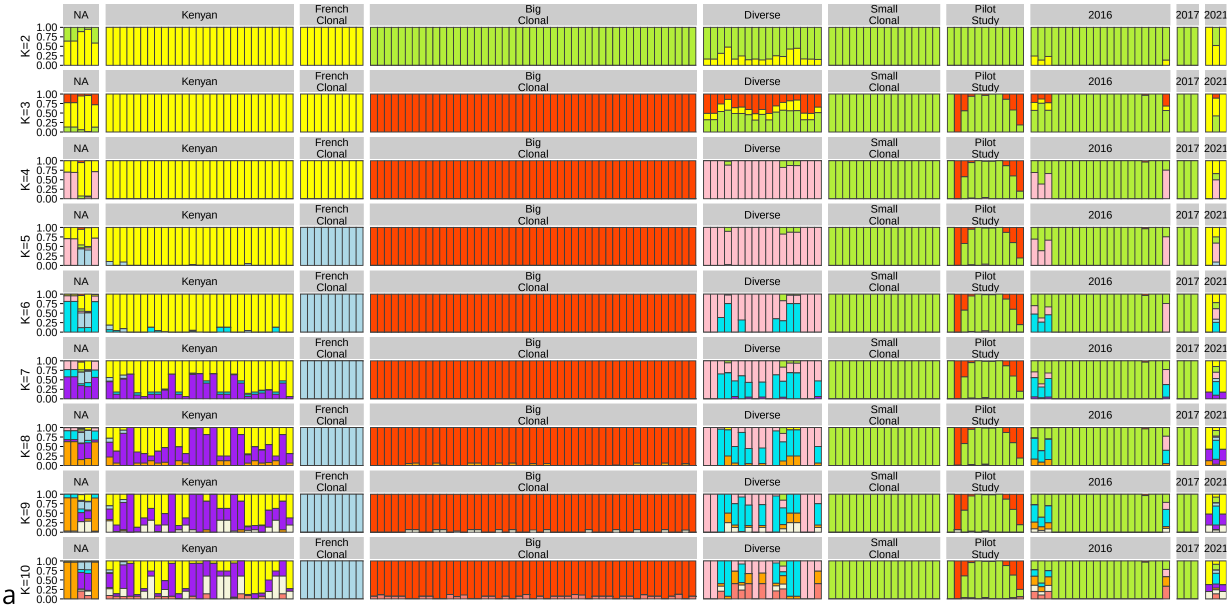


**Figure S4.** Admixture ancestry coefficients in K from 2 to 10 without pooled samples. Metagenmic samples are indicated by the year of sampling or pilot study and WGS Isolates are indicated by their lineage name

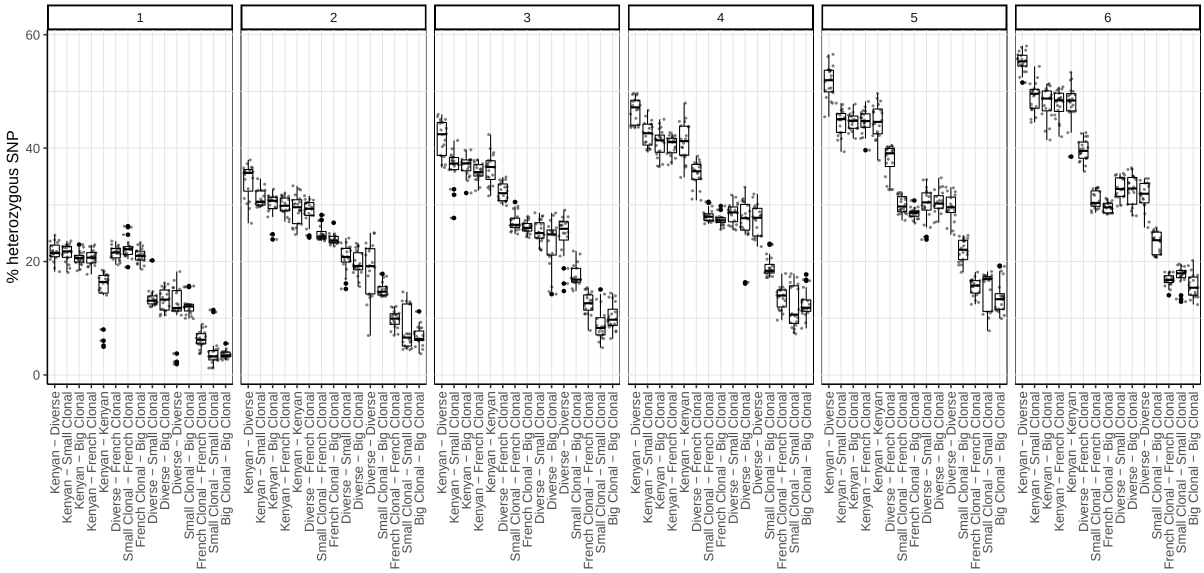


a

**Figure S5.** Percentage of heterozygous SNP in the artificial pools. Numbers indicate the number of samples per cluster included in the artificial pool (from 1 to 6)

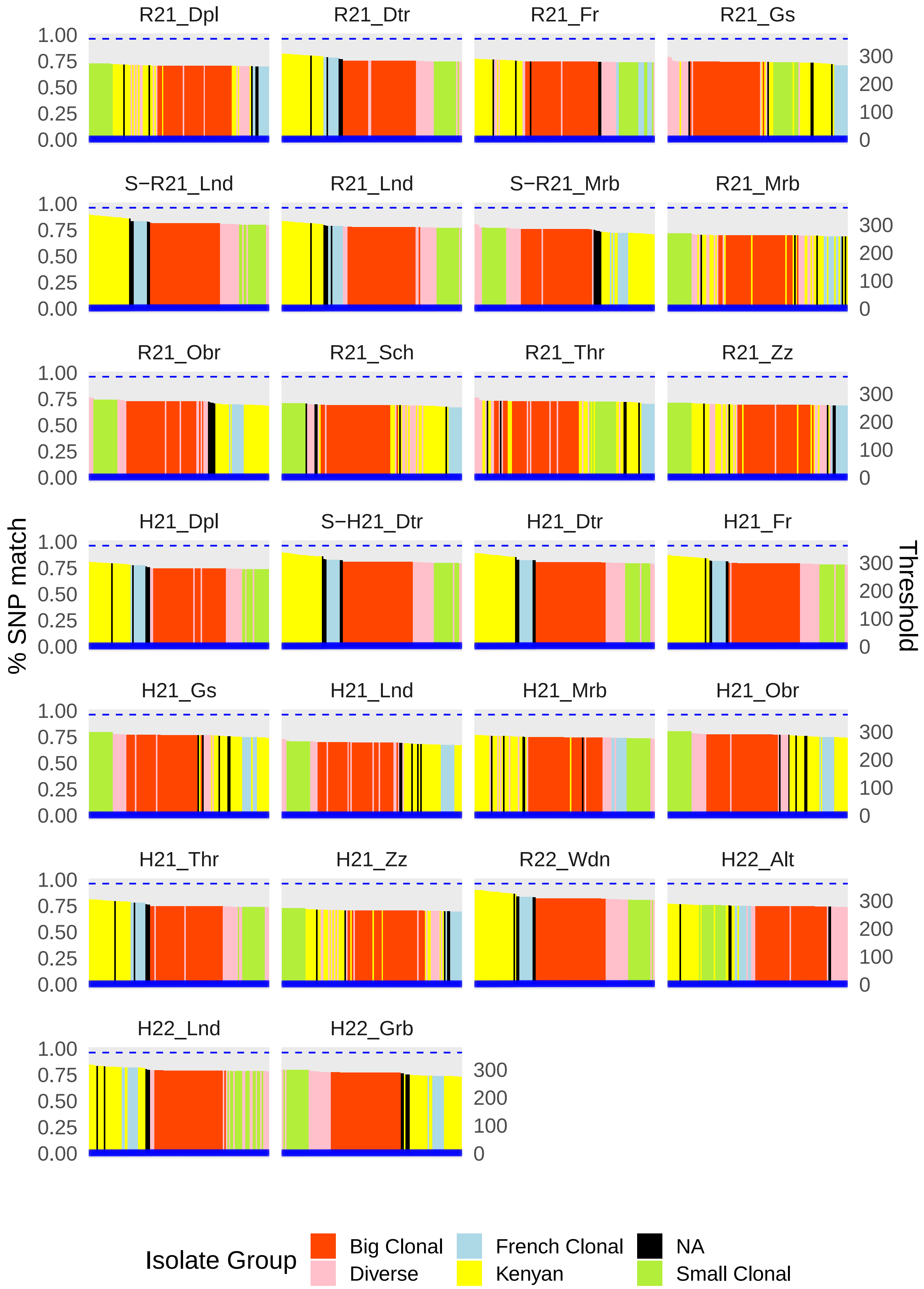


a

**Figure S6.** SNPmatch. Left y-axis presents the percentage of SNPs matching to the reference panel of WGS isolates (x-axis) in metagenomic samples with good coverage from 2021 and 2022. In blue and in the right y-axis, the likelihood ratio. Horizontal dashed blue line indicates the 3.841 treshold of significance.

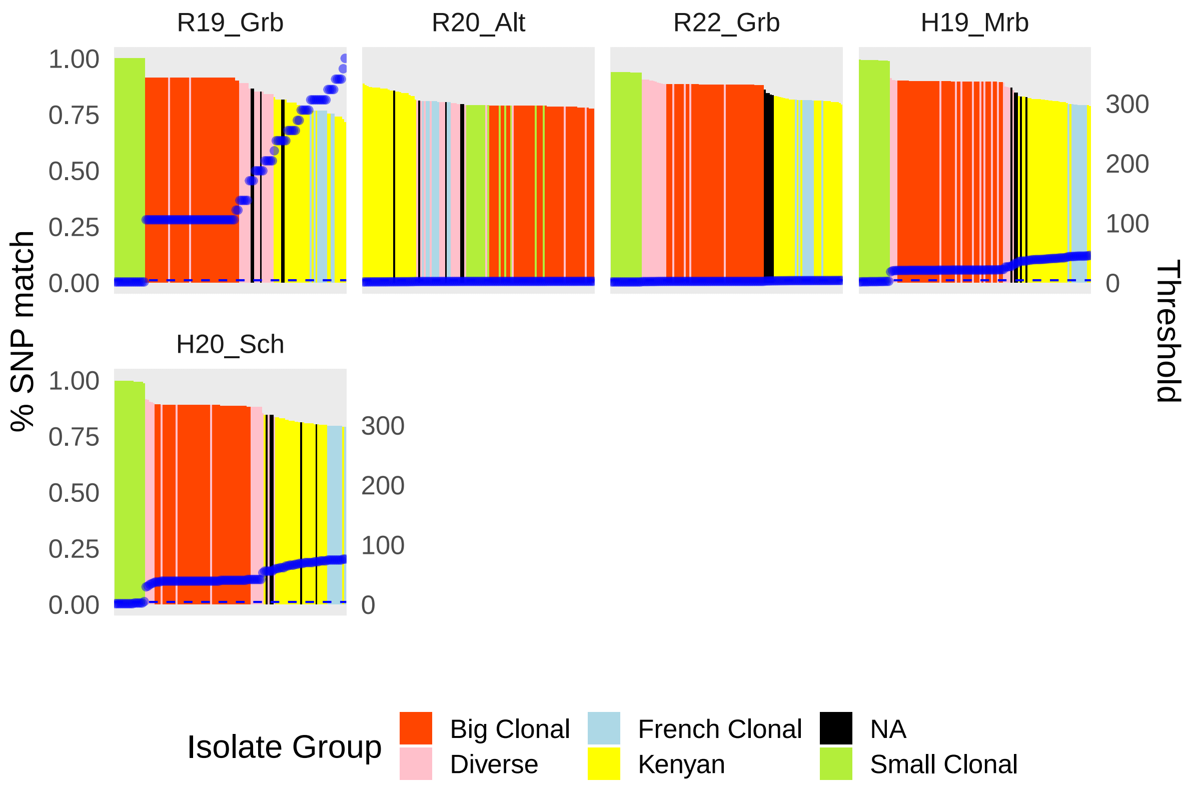


a

**Figure S7.** SNPmatch. Left y-axis presents the percentage of SNPs matching to the reference panel of WGS isolates (x-axis) in low-covered recovered metagenomic samples. In blue and in the right y-axis, the likelihood ratio. Horizontal dashed blue line indicates the 3.841 treshold of significance.

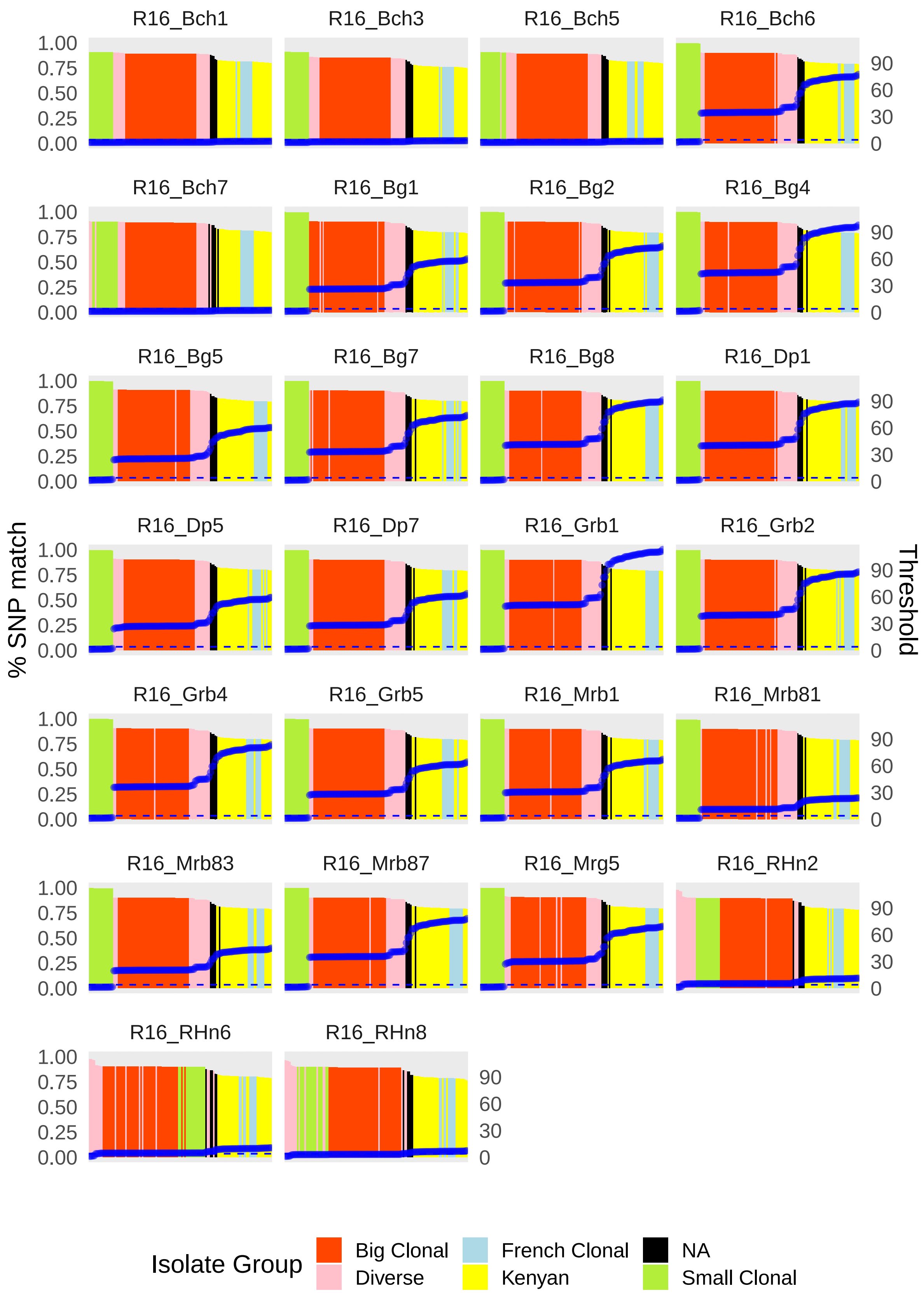


a

**Figure S8.** SNPmatch. Left y-axis presents the percentage of SNPs matching to the reference panel of WGS isolates (x-axis) in metagenomic samples from 2016 and 2017. In blue and in the right y-axis, the likelihood ratio. Horizontal dashed blue line indicates the 3.841 treshold of significance.

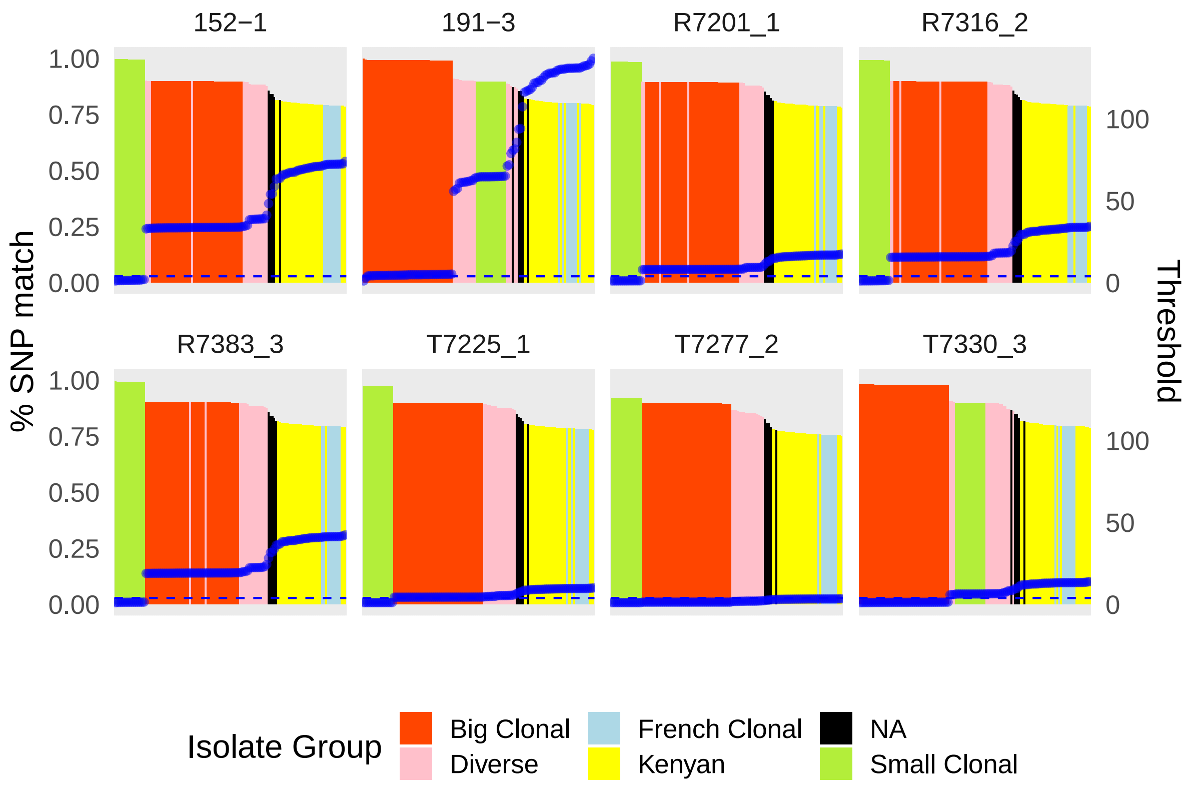


a

**Figure S9.** SNPmatch. Left y-axis presents the percentage of SNPs matching to the reference panel of WGS isolates (x-axis) in metagenomic samples from the pilot study. In blue and in the right y-axis, the likelihood ratio. Horizontal dashed blue line indicates the 3.841 treshold of significance.

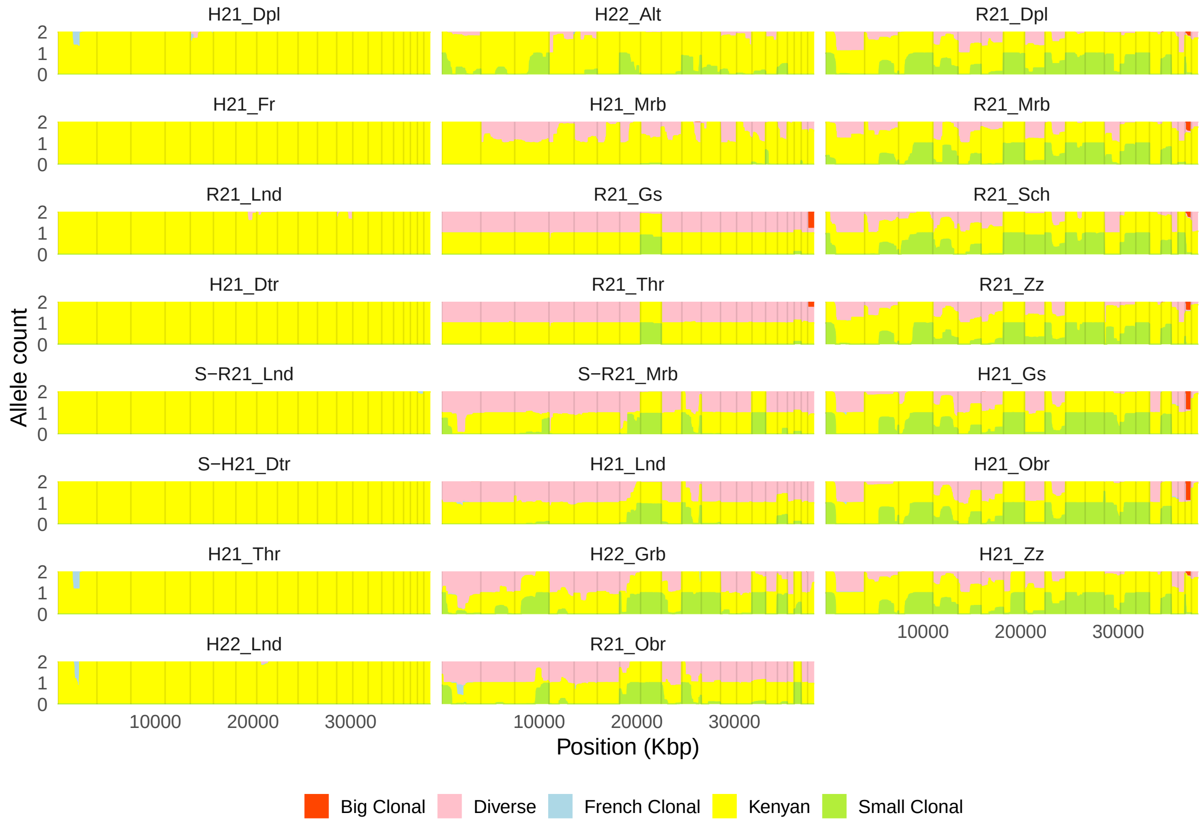


a

**Figure S10.** Local ancestry inference with ELAI in samples from 2021 and 2022. Plots indicate along the genome, the source population for each of the two alleles.

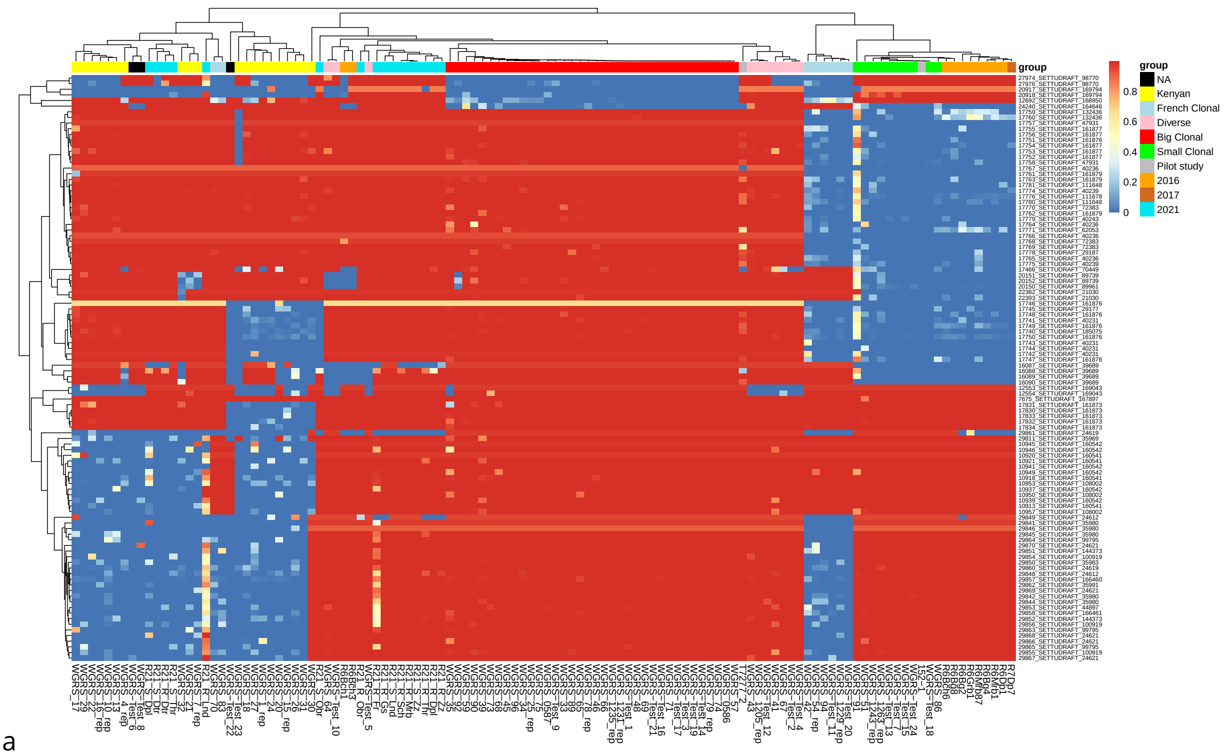


a

**Figure S11.** Hierarchical clustering of the samples with at least 10X mean coverage acording to the coverage of the CDS regions of a structured genes between European and Kenyan lineages. On top, cluster of the samples, on the left, cluster of the CDS sequences of each gene (SETTUDRAFT). WGS samples indicated by their lineage, metagenomic samples indicated by the sampling year

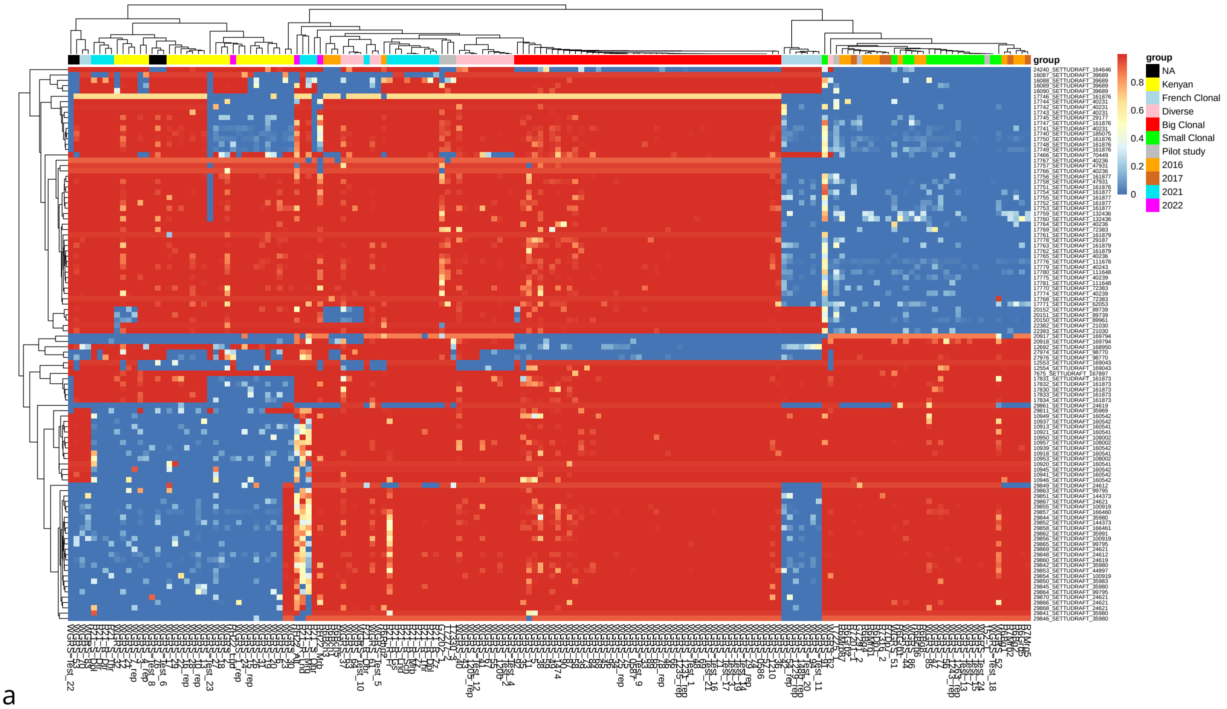


a

**Figure S12.** Hierarchical clustering of the samples with at least 4X mean coverage acording to the coverage of the CDS regions of a structured genes between European and Kenyan lineages. On top, cluster of the samples, on the left, cluster of the CDS sequences of each gene (SETTUDRAFT). WGS samples indicated by their lineage, metagenomic samples indicated by the sampling year

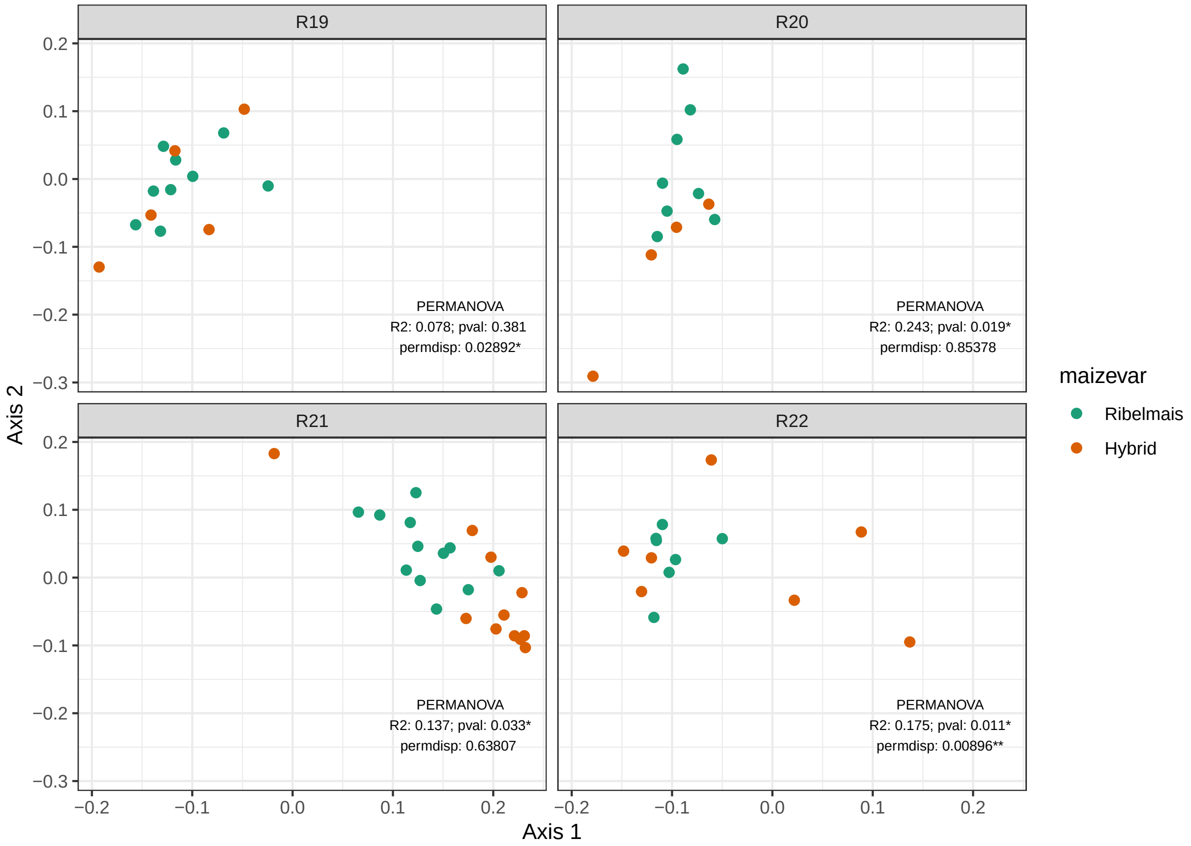


a

**Figure S13.** Principal Coordinate Analysis of the Bray-Curtis distance of phyllobiome composition for each year and PERMANOVA on the maize variety.
